# Broad transcriptional dysregulation of brain and choroid plexus cell types with COVID-19

**DOI:** 10.1101/2020.10.22.349415

**Authors:** Andrew C. Yang, Fabian Kern, Patricia M. Losada, Christina A. Maat, Georges Schmartz, Tobias Fehlmann, Nicholas Schaum, Davis P. Lee, Kruti Calcuttawala, Ryan T. Vest, David Gate, Daniela Berdnik, M. Windy McNerney, Divya Channappa, Inma Cobos, Nicole Ludwig, Walter J. Schulz-Schaeffer, Andreas Keller, Tony Wyss-Coray

**Author notes:** These authors contributed equally (Andrew C. Yang, Fabian Kern).

## Abstract

Though SARS-CoV-2 primarily targets the respiratory system, it is increasingly appreciated that patients may suffer neurological symptoms of varied severity^1–3^. However, an unbiased understanding of the molecular processes across brain cell types that could contribute to these symptoms in COVID-19 patients is still missing. Here, we profile 47,678 droplet-based single-nucleus transcriptomes from the frontal cortex and choroid plexus across 10 non-viral, 4 COVID-19, and 1 influenza patient. We complement transcriptomic data with immunohistochemical staining for the presence of SARS-CoV-2. We find that all major cortex parenchymal and choroid plexus cell types are affected transcriptionally with COVID-19. This arises, in part, from SARS-CoV-2 infection of the cortical brain vasculature, meninges, and choroid plexus, stimulating increased inflammatory signaling into the brain. In parallel, peripheral immune cells infiltrate the brain, microglia activate programs mediating the phagocytosis of live neurons, and astrocytes dysregulate genes involved in neurotransmitter homeostasis. Among neurons, layer 2/3 excitatory neurons—evolutionarily expanded in humans^4^—show a specific downregulation of genes encoding major SNARE and synaptic vesicle components, predicting compromised synaptic transmission. These perturbations are not observed in terminal influenza. Many COVID-19 gene expression changes are shared with those in chronic brain disorders and reside in genetic variants associated with cognitive function, schizophrenia, and depression. Our findings and public dataset provide a molecular framework and new opportunities to understand COVID-19 related neurological disease.

---

Severe Acute Respiratory Syndrome Coronavirus 2 (SARS-CoV-2), the etiologic agent of the COVID-19 global pandemic, has as of October 2020 infected over 40 million people and caused over a million deaths worldwide (worldometers.info/coronavirus). Though SARS-CoV-2 primarily targets the respiratory system, it is increasingly appreciated that COVID-19 patients can suffer neurological and psychiatric symptoms of varied severity, depending on SARS-CoV-2 mechanisms as well as other variables, such as co-morbidities and clinical care^1–3,5^. Central nervous system (CNS) symptoms range from loss of smell and headache to memory loss, encephalitis, stroke, difficulty concentrating, and persistent fatigue^1,6–8^. Symptoms are especially frequent and prominent in the over 20% of COVID-19 patients that require hospitalization^1,9–11^. With lingering CNS symptoms documented in Severe Acute Respiratory Syndrome (SARS) and Middle East Respiratory Syndrome (MERS) survivors^1,12–14^, it remains to be seen whether such chronic symptoms will emerge in COVID-19 survivors.

Cellular and molecular approaches are required to understand the neurological changes that may contribute to symptoms reported in COVID-19 patients. Fundamentally, pathology can arise from the direct infection of neurons and glia or indirectly from peripheral infection and its attendant immune response^15,16^. Viral RNA has been detected in the olfactory mucosa and cerebrospinal fluid (CSF) in some cases, but the consequences for brain cells remain unclear^17–20^. Several reports using elegant *in vitro* culture systems, such as neuronal organoids, have reached conflicting conclusions regarding the potential for SARS-CoV-2 neuroinvasion^21–26^. The physiological relevance of such cultured systems, from the viral titers applied to their fidelity to actual patient brains, also remains unclear^23,27,28^. Missing so far is an unbiased view of the molecular and cellular perturbations across brain cell types in COVID-19 patients. This is in part because high-quality human brain tissue from patients for such studies is largely inaccessible^21,23,24^. Furthermore, several brain regions highly relevant to COVID-19, such as the choroid plexus^23,24^, have not yet been successfully profiled in humans at single-cell resolution, in health or disease^29^.

Here, we profile 47,678 droplet-based single-nucleus transcriptomes from the frontal cortex and choroid plexus across 10 non-viral, 4 COVID-19, and 1 influenza patient (Fig. 1a). We complement transcriptomic data with immunohistochemical staining for the presence of SARS-CoV-2. These data—which includes the first, to our knowledge, single-cell view of neurological changes in COVID-19 and of the human choroid plexus in general—provide a molecular framework for how SARS-CoV-2 affects the brain without requiring direct neuroinvasion; reveals the major pathways and cell types perturbed; and highlights associations with chronic CNS diseases potentially pertinent to COVID-19 survivors. We created an interactive data browser (https://twc-stanford.shinyapps.io/scRNA_Brain_COVID19), which will be updated as more patient samples become available, to provide a resource for researchers to further dissect the molecular mechanisms of SARS-CoV-2’s impacts on the brain.

**Figure 1.**
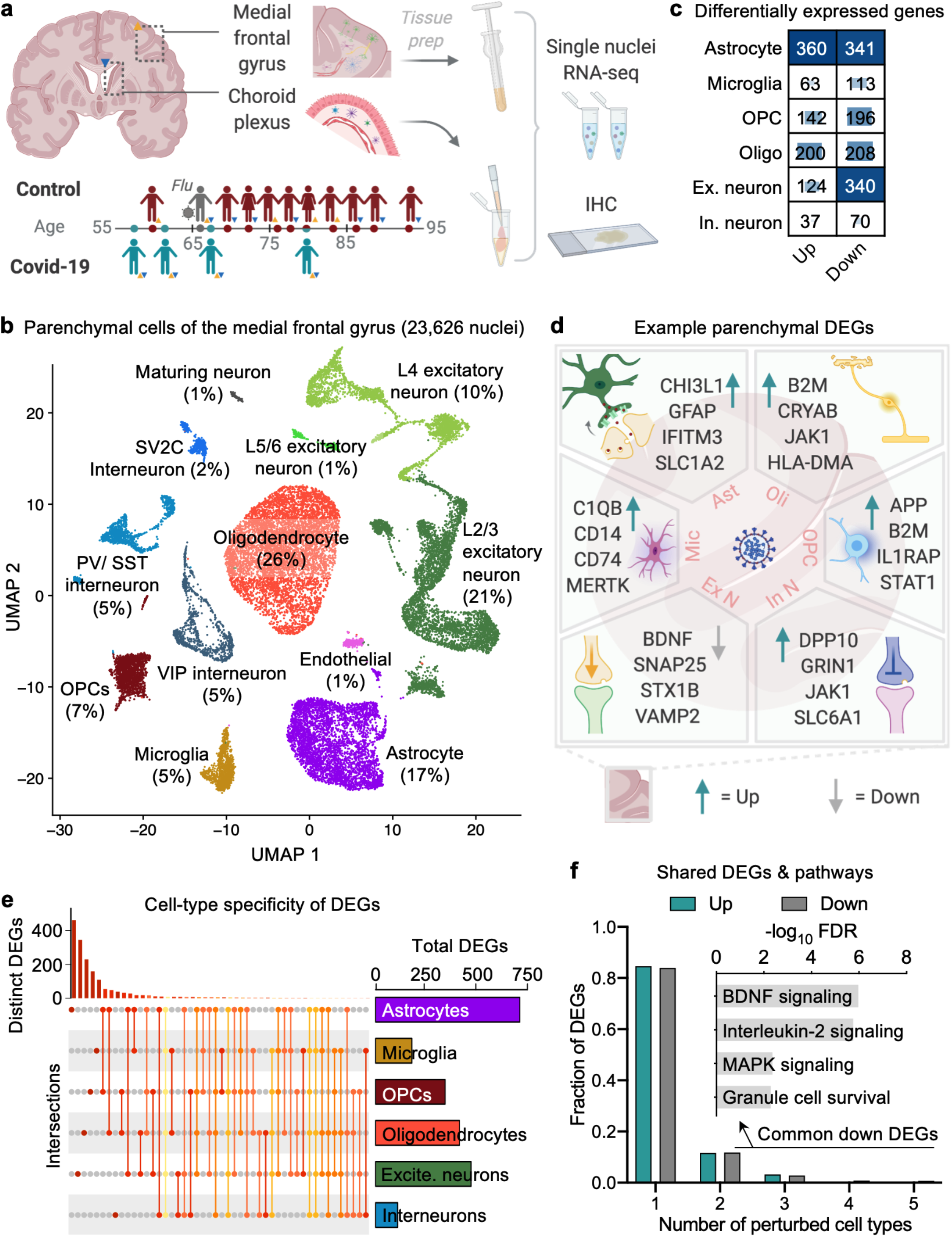
Cell type-specific gene expression changes in the brain of COVID-19 patients. **a**, Study design. Colored triangles denote brain regions studied for each patient. **b**, Uniform Manifold Approximation and Projection (UMAP) of 23,626 nuclei captured from the human medial frontal cortex, colored by cell type and labeled with percent of total nuclei. Note, that as in prior reports^32,34,35^, the ‘Endothelial’ cluster also exhibits vascular mural cell markers. **c**, Differentially expressed gene (DEG) counts for each cell type (MAST with default thresholds, FDR < 0.05, Log (fold change) > 0.25 (absolute value)). The intensity of the blue color and the size of the squares are proportional to entry values. **d**, Example differentially expressed genes (DEGs) in COVID-19: excitatory (Ex N) and inhibitory (In N) neurons, astrocytes (Ast), oligodendrocytes (Oli), oligodendrocyte precursor cells (OPC), and microglia (Mic). **e**, Matrix layout for intersections of DEGs shared across and specific to each cell type. Circles in the matrix indicate sets that are part of the intersection, showing that most DEGs are cell type-specific. **f**, Fraction of total up- and downregulated genes (y-axis) as a function of the total number of cell types in which the differential expression occurs. Biological pathways associated with genes downregulated in COVID-19 that are common to ≥2 cell types are shown (*n* = 161 genes, hypergeometric test, FDR correction).

## Cell type-specific gene expression changes in the COVID-19 brain

We aimed to gain insight into the cell type–specific transcriptomic effects of COVID-19 on the brain by performing unbiased single-nucleus RNA sequencing (snRNA-seq) of 15 fresh-frozen postmortem medial frontal cortex and lateral ventricle choroid plexus samples from 10 non-viral, 4 COVID-19, and 1 influenza patient of similar clinical course as the COVID-19 patients (Extended Data Table 1). Our sample sizes were similar to those reported in previous snRNA-seq studies^30–32^. Samples in the control and COVID-19 groups were between 58 and 91 years old and matched for tissue dissection area and RNA quality (Extended Data Fig. 1, Extended Data Table 1). Cause of death for COVID-19 and influenza patients was consistently interstitial pneumonia after more than two weeks of mechanical ventilation. Two COVID-19 patients experienced objective neurological symptoms during care (Extended Data Table 1). Samples were not confounded by technical or batch artifacts (Extended Data Fig. 2).

To begin, we generated 23,626 single-nuclei gene expression profiles from the medial frontal cortex (4 non-viral, 4 COVID-19, 1 influenza)—and detected a median of 1,486 genes per nucleus, a yield that is in line with recent snRNA-seq studies (Fig. 1b, Extended Data Fig. 1)^31–34^. We performed unbiased clustering of nuclear profiles to identify and annotate 8 cell types, including subtypes of excitatory neurons and interneurons. Each annotated cell type expressed previously established marker genes (Extended Data Fig. 3): excitatory neurons (e.g., marked by *CBLN2* and *NRGN*), inhibitory neurons (*GAD1*), astrocytes (*AQP4*), oligodendrocytes (*MBP* and *ST18*), microglia (*DOCK8*), oligodendrocyte progenitor cells (*VCAN*), maturing neurons (*DACH1*), and endothelial cells (*LEPR*)—though this latter population also contains mixed markers for vascular mural pericytes and smooth muscle cells, as in prior reports^32,34,35^. Overall, the cell type identities, markers, and proportions match prior snRNA-seq data from adult human cortex (Extended data Fig. 1–4, Extended Data Table 2)^31–34^.

We compared levels of gene expression in cells isolated from COVID-19 versus non-viral control individuals by cell type and identified 2,194 unique differentially expressed genes (DEGs) that implicated all major cell types (Extended Data Table 3). As recently reported in Alzheimer’s disease (AD)^34^, neurons exhibited a strong signature of repression—73% of DEGs in excitatory neurons and 65% in interneurons were downregulated—whereas DEGs in glial oligodendrocytes, OPCs, astrocytes, and microglia were more balanced (36-51%, Fig. 1c-d). The number of DEGs was greatest for astrocytes, excitatory neurons, and oligodendrocytes, partially reflecting increased power in these higher-abundance cell types. The vast majority of DEGs were perturbed in only a single cell type (~84%, Fig. 1e-f, Extended Data Fig. 5a), further indicating a cell type-specific CNS response to COVID-19. However, the minority of shared DEGs clustered into meaningful pathways: for example, commonly downregulated DEGs implicated impairments in brain-derived neurotrophic factor (BDNF) and interleukin-2 (IL-2) signaling (Fig. 1f), both critical for homeostatic neuronal function^36–40^. Overall, these results indicate that all major brain parenchymal cell types are affected at the transcriptional level by COVID-19, and that single-cell-level resolution is critical because changes in gene expression— including directionality—are dependent on cell type.

## SARS-CoV-2 accumulation in barrier cells stimulates inflammatory signaling to the brain

The observed transcriptional response to COVID-19 in the brain could arise from direct neurotropism or indirect damage via peripheral inflammation^15,16^. We thus stained for the SARS-CoV-2 spike glycoprotein^20,41^ in medial frontal cortex and choroid plexus tissue immediately adjacent to tissue used in snRNA-seq. Intriguingly, we detected spike glycoprotein robustly in two COVID-19 patients (Fig. 2). Upon closer examination however, spike glycoprotein signal resided only within the barrier-forming cortical vasculature, meninges, and choroid plexus. Within the choroid plexus, signal appeared within the stromal compartment, though confident cell type assignment was not possible (Fig. 2c). Thus, unlike recent reports where SARS-CoV-2 was added directly to neuroglial organoids lacking brain-barrier cells^21–25^, we do not observe significant virus invasion in cortex parenchymal cells. On the other hand, SARS-CoV-2 infection of the CSF-barrier forming choroid plexus and meninges corroborates recent *in vitro, in silico*, CSF, and postmortem case studies suggesting CNS barrier cells can support productive infection at relatively low titers^5,23,24,42,43^. We note however that this absence of detectable SARS-CoV-2 neuroinvasion does not exclude its possibility. Likewise, it is possible that the other two COVID-19 patients sustain SARS-CoV-2 barrier cell infection undetected in the tissue sampled. Corroborating a recent study^5^, presence of SARS-CoV-2 in the CNS did not correlate with neurological symptoms (Extended Data Table 1). Together, these results suggest that brain-barrier cells may be particularly susceptible to SARS-CoV-2 infection, and that preferential accumulation of virus in the barriers may give rise to widespread cortical DEGs.

**Figure 2.**
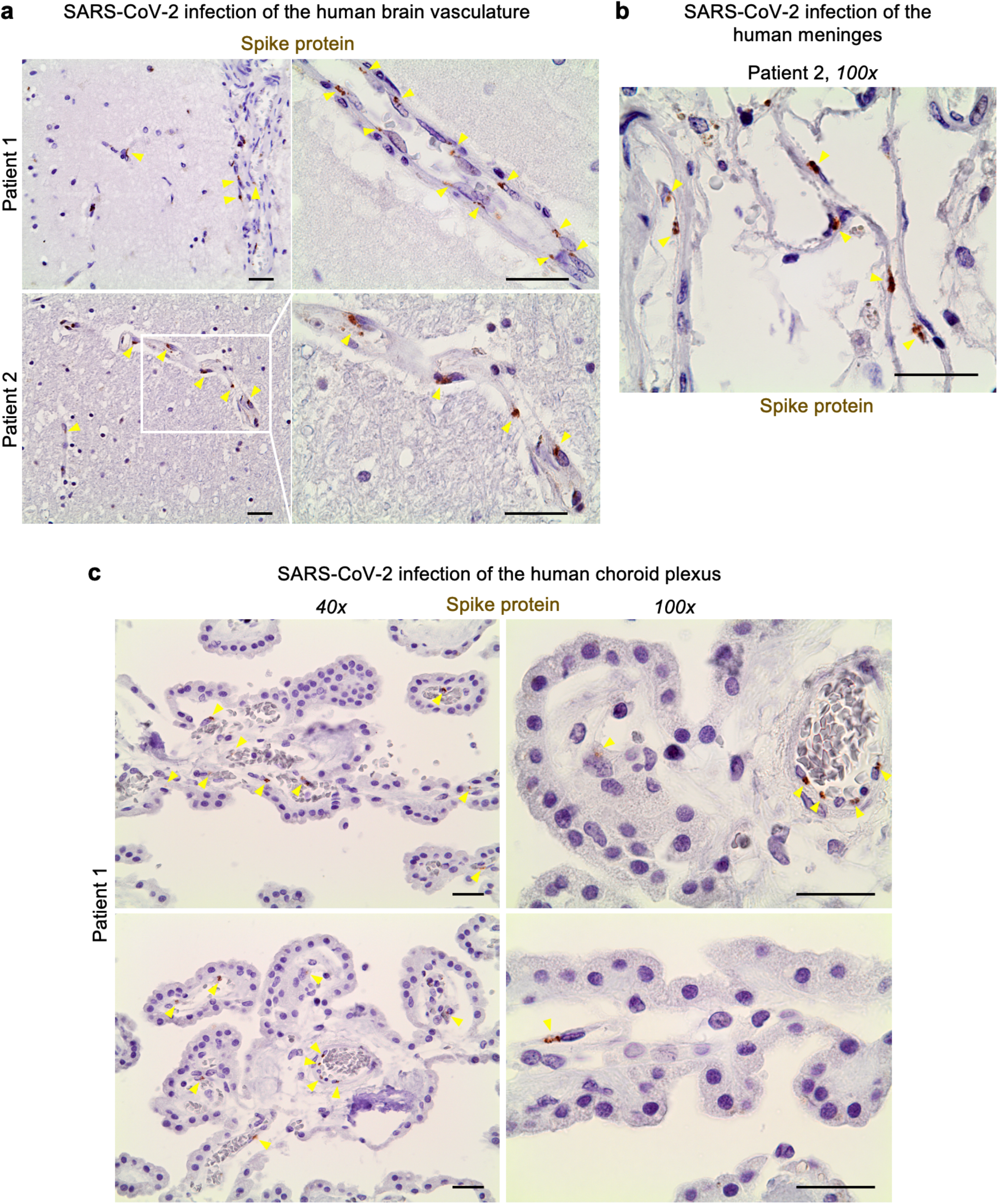
SARS-CoV-2 infection of blood-brain and -CSF barrier cells. **a**, SARS-CoV-2 spike glycoprotein (brown) detection in the frontal medial cortex of two COVID-19 patients in tissue immediately adjacent to that used for snRNA-seq. Hematoxylin counterstain (blue). Scale bar = 20 microns. **b**, As in (**a**) but for the meninges in Patient 2. Scale bar = 20 microns. **c**, As in (**a**) but for the choroid plexi immediately adjacent to tissue used for snRNA-seq, in Patient 1. Scale bar = 20 microns.

Because we detected SARS-CoV-2 spike glycoprotein in the blood-brain and blood-CSF-barrier cells, we sought to devise methods to profile these cells and better understand their relation to cortical transcriptional perturbations with COVID-19. However, conventional brain snRNA-seq preparation methods nearly universally deplete high-quality brain vascular nuclei from endothelial cells, pericytes, smooth muscle cells, and fibroblasts for currently unknown reasons^32–35^. At the same time, no snRNA-seq study exists on the human choroid plexus, in health or disease. This could, as we discovered, be because enzymatic and dounce homogenization techniques successfully used to process mouse tissue^29^ resulted in a near complete loss of genes in isolated human choroid nuclei. We thus generated a method involving successive gentle triturations in mild detergent (Methods), yielding 24,052 nuclei across 7 major epithelial, mesenchymal, immune, and glial cell types (6 non-viral, 4 COVID-19, 1 influenza; Fig. 3a, Extended Data Fig. 4, Extended Data Table 4).

**Figure 3.**
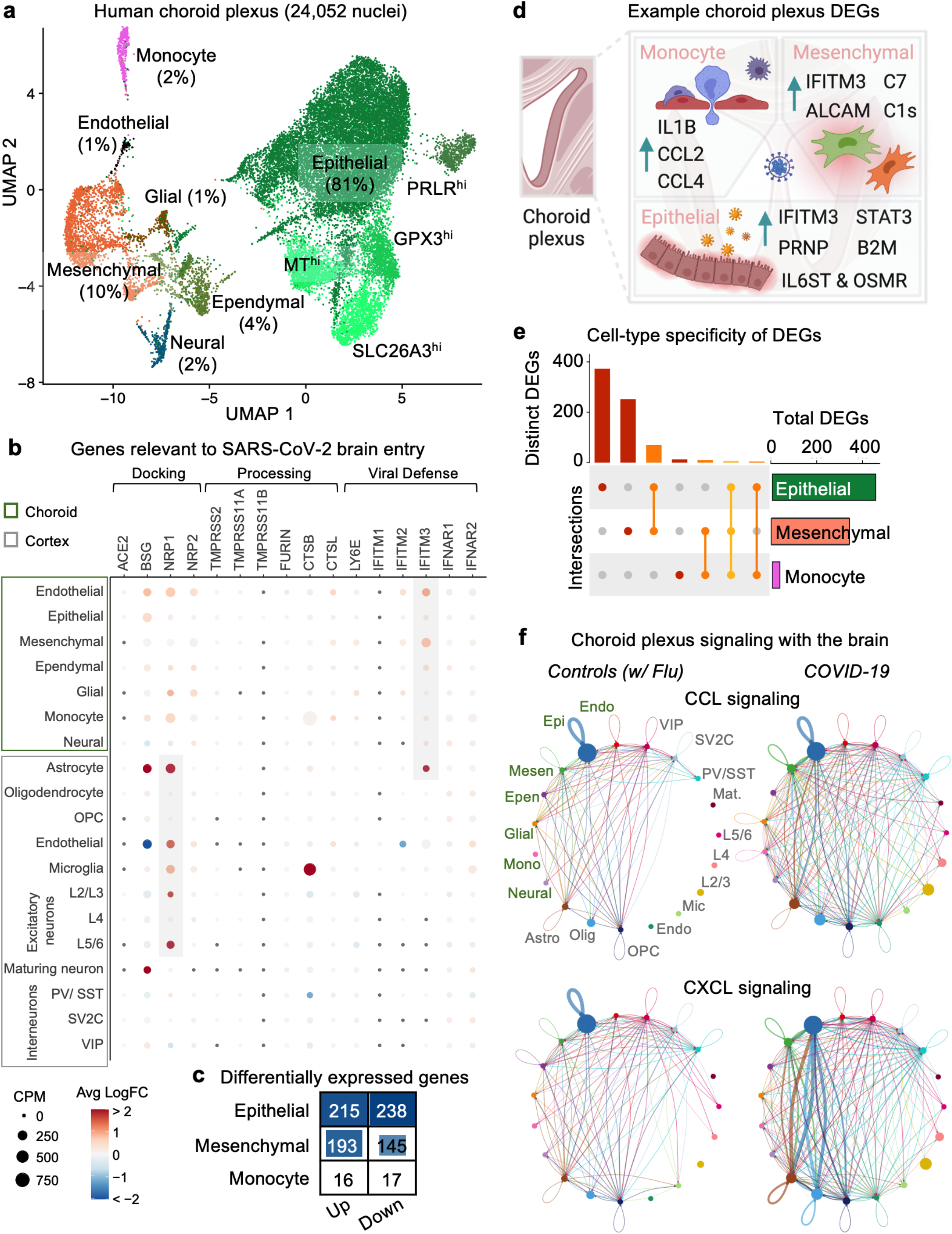
Viral defense and inflammatory signaling in blood-CSF barrier cells. **a**, UMAP of 24,052 nuclei captured from the human lateral choroid plexus, colored by cell type and labeled with percent of total nuclei. **b**, Expression profiles (CPM, circle size) and differential expression in COVID-19 patients (Log (fold change), color) for genes relevant to SARS-CoV-2 brain entry^16^. Grey indicates genes consistently upregulated in either brain-barrier cells (*IFITM3*) or cortex (*NRP1*). **c**, DEG counts for each cell type (MAST, FDR < 0.05, Log (fold change) > 0.25 (absolute value)). The intensity of the blue color and the size of the squares are proportional to entry values. **d**, Example DEGs in COVID-19 choroid plexus. **e**, Matrix layout for intersections of DEGs shared across and specific to each cell type. **f**, Circle plot showing the number of statistically significant intercellular signaling interactions for the CCL and CXCL family of molecules in controls (non-viral and influenza) compared to COVID-19 patients (permutation test, CellChat^61^).

Epithelial cells accounted for 81% of choroid nuclei profiled, and interestingly, subclustered based on expression of the receptor (PRLR) for prolactin—a hormone mediating neurogenesis^44,45^—, the glutathione peroxidase GPX3—which protects the CSF from hydroperoxides^46^—, and the chloride anion exchanger SLC26A3—which may maintain physiologic ion concentrations in the CSF^47^. Similar to brain vascular cells^16,48^, the choroid barrier and stromal epithelial, endothelial, mesenchymal, and ependymal cells robustly expressed the antiviral defense gene *IFITM3*. Intriguingly, we observed a broad upregulation of *IFITM3* across barrier cells in COVID-19 patients, consistent with immunohistochemical detection of SARS-CoV-2 infection (Fig. 3b). *IFITM3* serves as the first line of defense against viral infection^49^ and its upregulation is a marker of SARS-CoV-2 infection across publicly available datasets^50^. We also observed in the cortex parenchyma a consistent upregulation of the newly described viral entry receptor *NRP1*^41,51,52^ (Fig. 3b, Extended Data Fig. 6), which has been previously found upregulated in inflamed COVID-19 lung tissue^51,53^.

Overall, the DEGs detected in choroid epithelial cells, mesenchymal cells, and monocytes indicate strong CNS inflammation in COVID-19 (Fig. 3c-e, Extended Data Table 5). For example, we observed upregulation of the pro-inflammatory cytokine IL-1β, which has been reported sufficient to induce pathologic microglial and astrocyte activation^54,55^. The coincident upregulation of *IL-1β, CCL2*, and *CCL4* further suggests a weakened choroid barrier and an increasingly permissive environment for immune cell transmigration into the brain parenchyma^56–58^. The aged choroid plexus has recently been reported to send inflammatory signals to the brain, thereby activating glia and impairing cognitive function^59,60^. To assess if similar pro-inflammatory signaling mechanisms occur in COVID-19, we performed cell-cell communication analysis^61,62^. We observed a strong increase in choroid-to-cortex signaling across key inflammatory pathways, such as the CCL and CXCL family of chemokines (Fig. 3f, Extended Data Fig. 5b). Together, these results suggest that SARS-CoV-2 can infect and inflame brain-barrier cells, including the choroid plexus; and that this barrier—together with peripheral—inflammation is then relayed into the brain parenchyma.

## Immune cell infiltration and activation of neuronal engulfment programs in microglia

We sought to evaluate the predicted activation of brain microglia and migration of peripheral immune cells via enhanced inflammatory signaling from the periphery and brain-barrier cells. Intriguingly, we observed in the cortex a COVID-19 enriched subpopulation of microglia (Fig. 4a-b, Extended Data Fig. 7–8). This cluster was marked by expression of genes implicated in human microglial activation, such as *CD74, FTL*, and *FTH1* (Extended Data Table 6). These very genes have recently been described as difficult to detect by snRNA-seq^63^, further underscoring strong COVID-19 microglial activation. An over-represented fraction of COVID-19 microglia cluster marker genes overlap (*P* < 10^−25^, hypergeometric test) with human Alzheimer’s disease microglia cluster markers^34^ (Extended Data Fig. 7c). Monocle trajectory analysis^64^ found that the COVID-19 microglia cluster emerged from the parent homeostatic population—and that the expression of activation markers increased along the pseudotime route—, suggesting that these microglia emerged in response to an increasingly inflamed CNS environment (Fig. 4c, Extended Data Fig. 7a-b), consistent with enhanced brain-barrier inflammatory signaling. Our observations suggest that the COVID-19 subpopulation represents a distinct microglial state that shares features with, but is ultimately different from, microglial cell states previously reported in human neurodegenerative disease.

**Figure 4.**
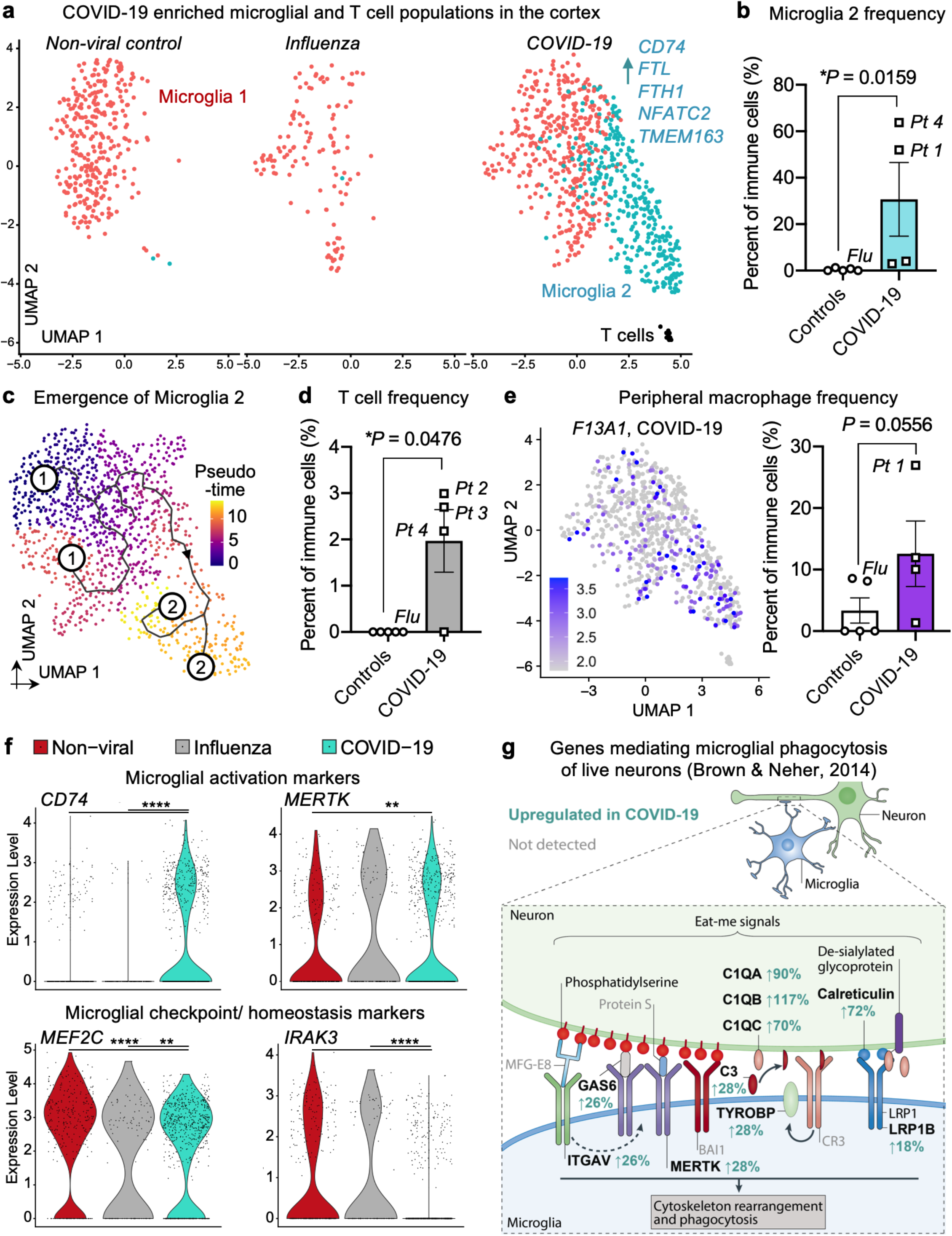
Immune cell infiltration and activation of microglial phagocytosis programs in COVID-19 brains. **a**, UMAP of 1,154 immune cells captured in the human frontal cortex, split by non-viral controls, influenza, and COVID-19 patients. Cells are colored by cell type subcluster. Example genes upregulated in Microglia cluster 2 are shown. **b**, Quantification of Microglia 2 population as a proportion of total immune cells (*n* = 5 controls, including influenza; *n* = 4 COVID-19, two-sided t-test; mean +/− s.e.m.). **c**, Monocle^64^ pseudotime trajectory plotting the emergence of Microglia cluster 2. Pseudotime is indicated in graded purple (high) to yellow (high). Numbers indicate Microglial cluster from (**a**). **d**, Quantification of the T-cell population as a proportion of total immune cells (*n* = 5 controls, including influenza; *n* = 4 COVID-19, two-sided t-test; mean +/− s.e.m.). **e**, Expression of the established peripheral macrophage-specific marker *F13A1*^65,66^. Expression (logCPM) is indicated in graded purple (left). Quantification of the peripheral macrophage population as a proportion of total immune cells (*n* = 5 controls, including influenza; *n* = 4 COVID-19, two-sided t-test; mean +/− s.e.m., right). **f,** Expression of established microglial activation^67–74^ and inflammation checkpoint^75–77^ molecules. Violin plots are centered around the median, with their shape representing cell distribution (\*\*\**P* < 0.001, \*\*\*\**P* < 0.00001, MAST). **g**, Upregulation of genes mediating the microglial phagocytosis of live neurons^81^. Bold indicates upregulated genes in COVID-19, grey not detected.

We next assessed the predicted increase in peripheral immune cell infiltration. We indeed found both an increase in T-cells (marked by *CD247* encoding the TCR CD3-zeta protein, Fig. 4d, Extended Data Fig. 8e) and an increase in peripheral macrophages (marked by *F13A1*^65,66^, Fig. 4e). Microglial activation and enhanced immune cell infiltration have been recently reported by IHC staining in a series of 43 COVID-19 brains, with the degree of infiltration and glial activation not clearly correlated with neurological symptoms or the presence of detectable virus in the brain^5^. Our results largely corroborate those findings via an orthogonal snRNA-seq approach (Fig. 4a-e). Both the formation of a distinct microglial cluster and the enhanced infiltration of peripheral T-cells and macrophages did not occur in terminal influenza, suggesting a COVID-19 specific—or at least not a pan-viral—CNS response.

Because we collected sufficient microglia for DEG analysis, we sought a clearer understanding of the functional ramifications of their COVID-19 activation. As expected, established markers of microglial activation previously linked to pathologic functions such as *CD74* and *MERTK* were strongly upregulated^67–74^. We observed a concomitant loss of the immune checkpoint molecules *MEF2C* and *IRAK3* that restrain runaway inflammation^75–77^. Recently, increased microglial phagocytosis of live neurons—termed phagoptosis—has been demonstrated to be an important mechanism driving various neurodegenerative diseases^71,78–81^. Remarkably, nearly every gene reported^81^ to mediate microglial phagocytosis of live neurons is strongly upregulated in COVID-19 microglia (Fig. 4g). These include the genes encoding complement proteins C3, all three components of C1Q, and the pro-phagocytic receptors MERTK and VNR. Together, these data reveal in the COVID-19 brain the infiltration of immune cells and strong activation of a pathologic microglial phagocytosis program predicted to prematurely engulf and prune live neurons.

## Potential synaptic dysfunction specifically in layer 2/3 excitatory neurons

The global induction of microglial phagoptosis programs prompted us to look into other potential signatures of neuronal compromise. Astrocytes play a dual role in both forming the glia limitans barrier^82,83^ and supporting homeostatic neuronal function^84–86^ (Fig. 5a). Given SARS-CoV-2 infection of barrier cells, we thus wondered whether astrocytes would exhibit similar hallmarks of inflammation with concomitant consequences for neuronal health. We indeed observed hallmarks of astrogliosis^87^ and viral defense^50^ in upregulated *GFAP* and *IFITM3* expression, as well as neurotoxicity in the secreted factor CHI3L1^88–90^ (Fig. 3b, Fig. 5a-d). Forming a tripartite synapse with neurons, astrocytes finely balance the levels of synaptic neurotransmitters like glutamate via the dedicated transporters EEAT1 (*SLC1A3*) and EEAT2 (*SLC1A2*)^91–93^. Dysregulated glutamate transporter expression causes neuronal dysfunction: downregulation results in excess glutamate excitotoxicity^94,95^ and extrasynaptic spillover^96,97^; whereas upregulation impairs long-term potentiation (LTP) and synaptic plasticity^98–101^. We observed dysregulated overexpression of genes predicting diminished glutamate signaling across synapses: for example, the overexpression of both glutamate transporters (*SLC1A2* and *SLC1A3*, Fig. 5a, e) as well as the major potassium uptake channel Kir4.1 (*KCNJ10*), which reduces neuronal excitability by lowering extracellular potassium, glutamate, and BDNF^102^.

**Figure 5.**
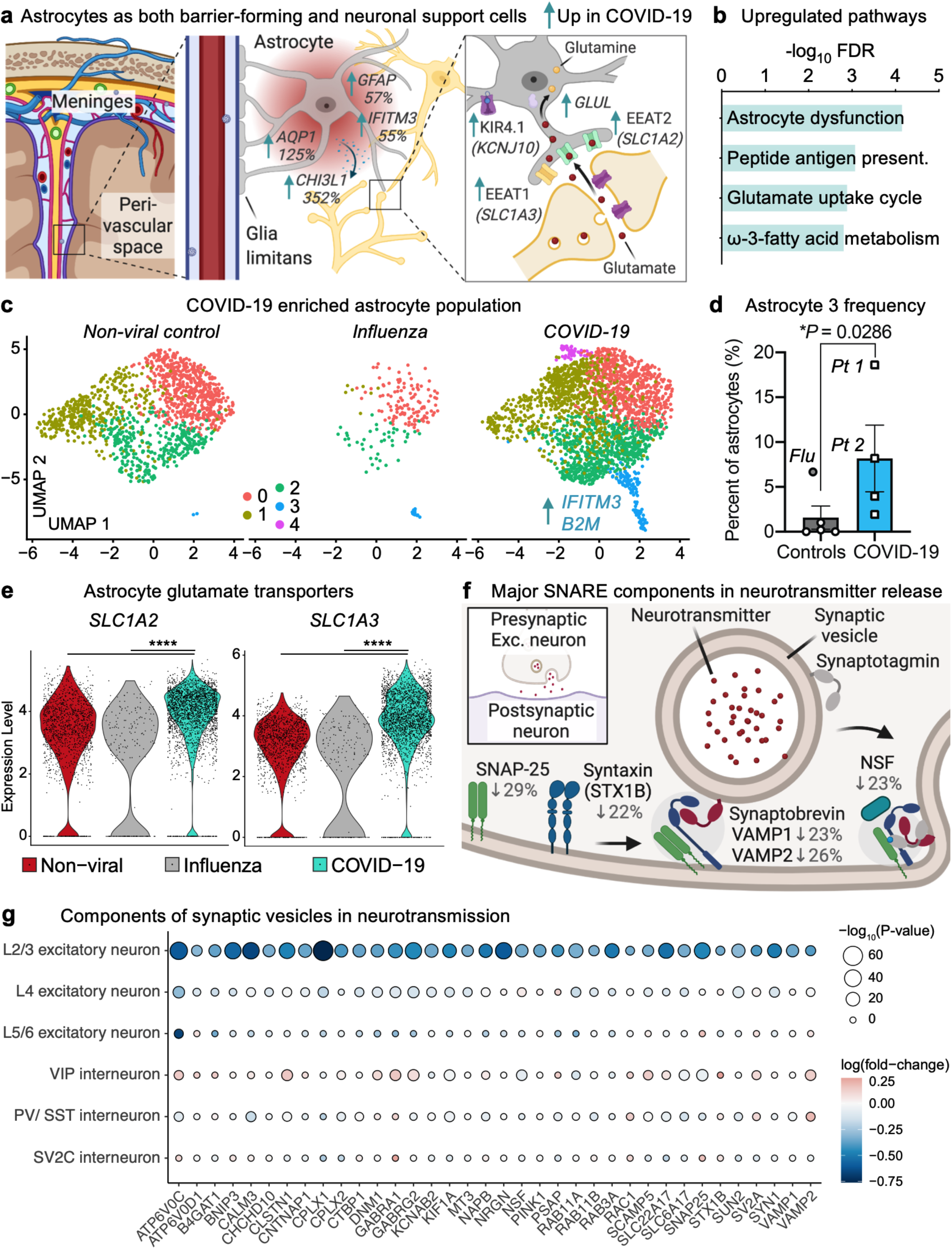
Impaired astrocytic support and excitatory neurotransmission in COVID-19 cortex. **a**, Upregulation of inflammatory and dysregulation of homeostatic genes in COVID-19 astrocytes (*n* = 3,967). **b**, Enriched biological pathways (Elsevier Pathway Collection, Enrichr) amongst upregulated genes in COVID-19 astrocytes. Enrichment is based on FDR-corrected cumulative hypergeometric *P* values, with a *P* value-thresholded DEG gene-marker list (Log (fold change) > 0.25 (absolute value), adjusted *P* value (Bonferroni correction) < 0.05, MAST). **c**, UMAP of 3,967 astrocytes captured in the human frontal cortex, split by non-viral controls, influenza, and COVID-19 patients. Cells are colored by cell type subcluster. Example genes upregulated in Astrocyte cluster 3 are shown. **d**, Quantification of Astrocyte 3 population as a proportion of all astrocytes (*n* = 5 controls, including influenza; *n* = 4 COVID-19, two-sided t-test excluding influenza patient; mean +/− s.e.m.). **e**, Expression of homeostatic astrocyte glutamate transporter genes encoding EEAT2 (*SLC1A2*) and EEAT1 (*SLC1A3*). Violin plots are centered around the median, with their shape representing cell distribution (\*\*\*\**p* < 0.0001, MAST). **f**, Downregulation of major SNARE components in COVID-19 excitatory neurons (*n* = 7,680 excitatory neurons). **g**, Heatmap showing downregulation of synaptic vesicle components specifically in COVID-19 L2/3 excitatory neurons (*n* = 7,680 excitatory neurons; *n* = 4,948 L2/3 excitatory neurons, Log (fold change) > 0.25 (absolute value), adjusted *P* value (Bonferroni correction) < 0.05, MAST).

We next sought to identify the neuronal subtypes affected in COVID-19. Enrichment analysis implicated potential impairments in synaptic transmission specifically in excitatory neurons (Extended Data Fig. 9). Among COVID-19 patients, those with objective neurological symptoms or detectable virus did not exhibit further deficits in the expression of genes mediating synaptic transmission (Extended Data Fig. 9c, Extended Data Table 1). Glutamate release at synapses is critical for fast and faithful communication between neurons for information processing in sub-second timeframes^103–105^. Glutamate is stored in readily releasable pools of vesicles within presynaptic terminals. During synaptic transmission, SNARE proteins play an indispensable role in vesicle docking, priming, fusion, and synchronization of glutamate release into the synaptic cleft^103,104^. Downregulation of the mRNA encoding SNARE components has been found in various neurological diseases^106–109^, and is sufficient in experimental models to impair both vesicle processing and overall neuronal viability^110–114^. We observed in COVID-19 excitatory neurons a consistent downregulation of major SNARE components (Fig. 5f). For example, genes encoding both v-SNARE synaptobrevins (*VAMP1, VAMP2*) necessary for calcium-dependent glutamate release^115,116^ decreased. Likewise, expression diminished for the partner t-SNARE proteins syntaxin 1B (*STX1B*) and SNAP-25, also necessary^117,118^ for glutamate release.

The human neocortex is composed of six layers of interconnected excitatory and inhibitory neurons, with each layer both highly specialized locally and integrated globally. Of these six layers, layers 2 and 3 are disproportionately thickened during the evolution of the human brain, playing a key role in higher brain functions such as cognition, learning, memory, creativity, and abstraction^4,119,120^. Though we captured neurons from all cortical layers, we found that gene expression changes linked to synaptic deficits are specific to only these layer 2/3 excitatory neurons (Fig. 5g). L2/3 excitatory neurons already exhibit sparse action potential firing to generate a simple and reliable neural code for associative learning^120^. Thus, this neuronal population may be particularly sensitive to the predicted deficits in neurotransmission with COVID-19. Together, these data suggest potential impairments in L2/3 neuron synaptic transmission, plasticity, and LTP in COVID-19, due to the dysregulation of genes mediating astrocyte glutamate homeostasis and genes encoding major synaptic vesicle components.

## Associations with chronic CNS disease

Reports of lingering CNS complications in COVID-19 survivors are emerging^121^. To provide a molecular rationale for these complications, we compared COVID-19 cell type-specific DEGs with those recently described by snRNA-seq in chronic CNS diseases, including Alzheimer’s disease^34^, multiple sclerosis^122^, Huntington’s disease^123^, and autism spectrum disorder^33^. Surprisingly, the overlap of DEGs between COVID-19 and these chronic CNS diseases was over-represented for each cell type (Fig. 6a, Extended Data Fig. 10, Extended Data Table 7). The degree of overlap was particularly high in astrocytes and excitatory neurons. For example, genes encoding the SNARE protein SNAP-25 and the neurotrophic factor BDNF were commonly downregulated in excitatory neurons in both COVID-19 and Alzheimer’s disease^34^. This over-representation of overlapping DEGs suggests existing CNS disease knowledge—and even treatment approaches—may be relevant to COVID-19 neuropathology.

**Figure 6.**
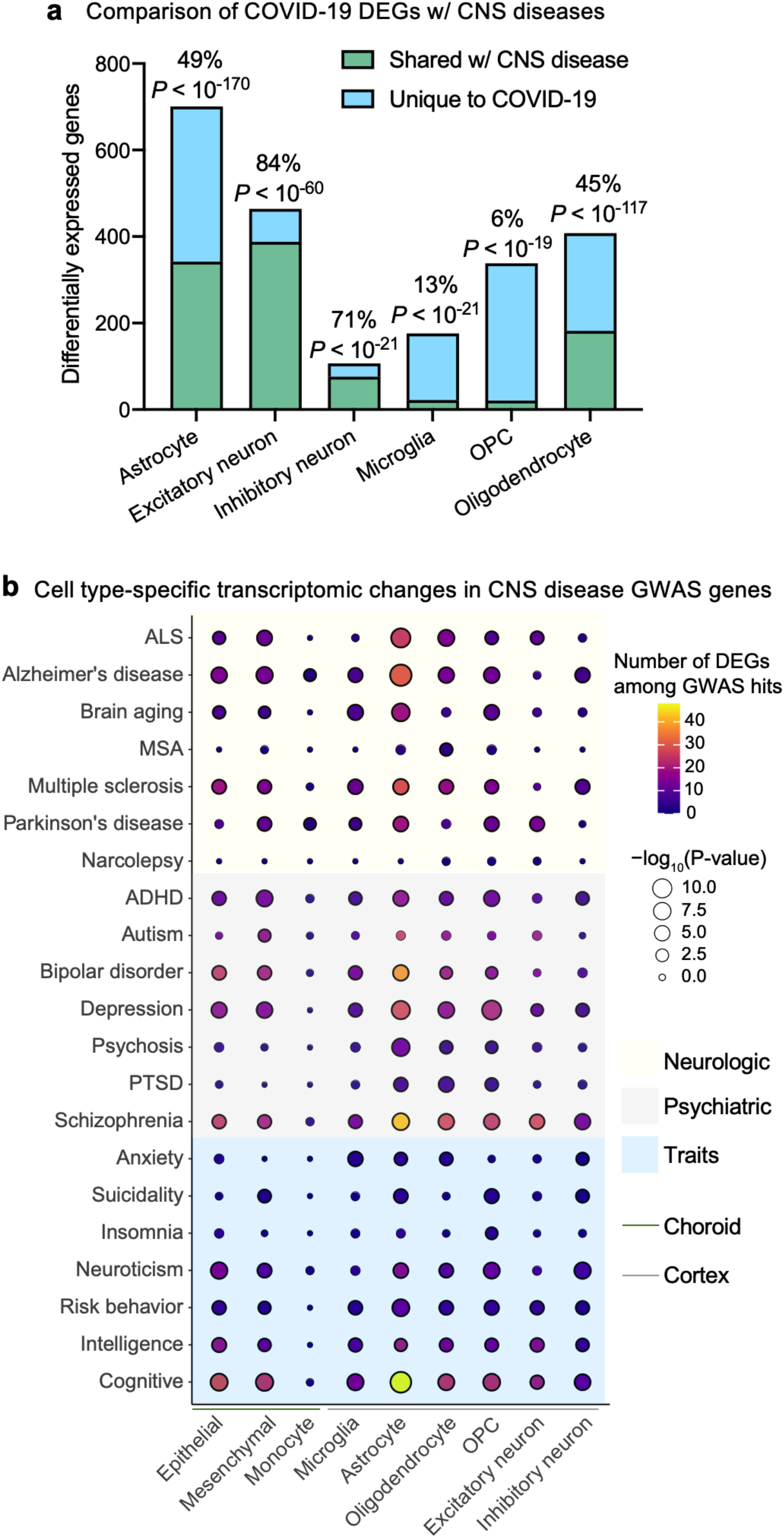
Association of COVID-19 gene expression changes with chronic CNS disease. **a**, Proportion of COVID-19 DEGs across cell types that are also DEGs in chronic CNS diseases, as published for Alzheimer’s Disease (AD)^34^, Autism spectrum disorder (ASD)^33^, Huntington’s disease (HD)^123^, and Multiple sclerosis (MS)^122^. Bars labeled with percent overlapping and over-representation *P* values (cumulative hypergeometric test). **b**, Heatmap showing the number of differentially expressed genes (DEGs) per cell type that are also GWAS risk variants across psychiatric and neurological diseases and traits from the GWAS catalog (NHGRI-EBI)^126^. Significance is based on FDR-corrected cumulative hypergeometric *P* values, with a *P* value-thresholded DEG list (Log (fold change) > 0.25 (absolute value), adjusted *P* value (Bonferroni correction) < 0.05, MAST). ALS: Amyotrophic lateral sclerosis, MSA: Multiple system atrophy, ADHD: Attention deficit hyperactivity disorder, and PTSD: Post-traumatic stress disorder.

Genome-wide association studies (GWAS) have shed insight into the molecular pathways involved in complex traits and diseases^124,125^. GWAS variants suggest an important role for a given gene through its loss or gain of function or expression. Thus, differential expression of GWAS genes in COVID-19 may signal a re-emergence of these trait and disease pathways^32^. To determine the enrichment of COVID-19 DEGs within genetic variants associated with complex traits and diseases in a cell type-specific fashion, we obtained GWAS summary statistics for neurological and psychiatric disorders and neurobehavior traits (Extended Data Table 8)^125–127^. We found a strong enrichment of DEGs residing within GWAS hits of various disorders and traits, especially in cognition, schizophrenia, and depression (Fig. 6b). These particular disorders have been associated with dysfunction in L2/3 neurons and astrocytes^128–135^. Together, these data suggest that COVID-19 may partially recapitulate established pathological processes in the brain and provide molecular hypotheses for emerging neurological complaints in COVID-19 survivors.

## Discussion

Recent snRNA-seq studies have elucidated the cell type-specific processes involved in several CNS diseases^32–35,122,123^. Here, by combining sequencing of 47,678 nuclei in both frontal cortex and choroid plexus along with IHC detection of SARS-CoV-2 in brain-barrier cells, we reveal several major neuropathological mechanisms in severe COVID-19. Based on gene expression patterns uncovered in this study, we propose a model where SARS-CoV-2 infected brain-barrier cells can relay and permit inflammatory signaling into the cortex parenchyma. In this inflammatory milieu, peripheral immune cells infiltrate the brain, microglia activate neuronal phagocytosis programs, and astrocyte homeostatic properties become dysregulated. Microglial activation is sufficient to induce a COVID-19 specific subpopulation. Together, these perturbations may converge on impaired neuronal function, with L2/3 excitatory neurons in particular exhibiting a transcriptomic signature of compromised glutamate transmission and information processing^4,119,120^.

There are, however, limitations to consider given the unique logistical circumstances of a pandemic. With most postmortem COVID-19 brain tissue immediately fixed^42^ or not handled with snRNA-seq studies in mind for safety and regulatory reasons, there is a lack of the high-quality tissue necessary for such studies^21,23,24^. This has precluded larger study cohorts that could be more representative of COVID-19 patients as a whole. For example, while we and a recent IHC-based study of 43 patient brains^5^ (the largest to date) do not find a clear correlation between detectable brain SARS-CoV-2 and glial inflammation or neurological symptoms, this may not be representative of all patients, especially those without pre-existing co-morbidities or who do not require ventilation^136^. Furthermore, the associations with chronic CNS disease may not be relevant in mild COVID-19 disease and may change over time in severe COVID-19 survivors. Long-term postmortem studies of survivors after the acute phase of disease are not currently possible to assess this. However, there is precedent for acute viral infections causing long-term inflammation and dysfunction predisposing neurodegenerative disease^137–140^.

Understanding the underlying mechanisms of how SARS-CoV-2 affects the brain may inform therapeutic approaches^121^. Most molecular studies to date on SARS-CoV-2 neurotropism have made powerful use of cultured organoids, though they have reached conflicting conclusions^21–26^. Here, in postmortem COVID-19 patient tissue, we find that the blood-brain and blood-CSF barrier cells typically lacking in organoid models are of high physiologic relevance in informing questions of neurotropism. By combining IHC with a new method to process human choroid plexi tissue for snRNA-seq, we propose that infection of brain-barrier cells can generate a depot of strong antiviral inflammation and compromise barrier function, combining to inflame the brain parenchyma. Such brain-barrier relay and permeability mechanisms have been found important in brain aging^59,141–144^. Thus, COVID-19’s impact on the brain, like many of its other properties, may follow a nuanced—but potentially therapeutically targetable—mechanism of action. We propose that the development of new techniques to study human brain-barrier cells will further help elucidate disease processes in COVID-19 and beyond.

The scale of the COVID-19 pandemic means thousands of people may suffer neurological symptoms, with some facing lifelong problems as a result. Our results nominate COVID-19 pathologic processes in microglia, astrocytes, and neurons not seen in a patient suffering terminal influenza infection. The particular vulnerability of layer 2/3 excitatory neurons in COVID-19 provides a consistent molecular hypothesis for emerging neurological complaints, from “brain fog” to memory loss to difficulty concentrating^11,145,146^. Several COVID-19 perturbations revealed here may be addressable with existing medications^147^. However, as is the case for recent human transcriptomic studies, additional research will be required to link the observed cell type-specific gene expression changes with neurological symptoms^32,33,35,122,123^. Nevertheless, these studies have served as important resources for understanding the molecular and cellular basis of disease and for formulating new lines of investigation. In summary, this work sheds insight into and provides new opportunities to explore the acute and lasting neurological consequences of COVID-19.

## Supporting information

ED Table 1 Patient Information

ED Table 2 Cortex Parenchyma Markers

ED Table 3 Cortex Parenchyma DEGs

ED Table 4 Choroid Plexus Markers

ED Table 5 Choroid Plexus DEGs

ED Table 6 Microglia COVID subcluster markers

ED Table 7 DEG overlap CNS disease

ED Table 8 CNS GWAS genes

## Acknowledgments

We thank N. Khoury, T. Iram, E. Tapp, O. Hahn, and other members of the Wyss-Coray and Keller labs for feedback and support, and H. Zhang and K. Dickey for laboratory management. This work was funded by the NOMIS Foundation (T.W.-C.), the National Institute on Aging (T32-AG0047126 to A.C.Y., 1RF1AG059694 to T.W.-C), Nan Fung Life Sciences (T.W.-C.), the Bertarelli Brain Rejuvenation Sequencing Cluster (an initiative of the Stanford Wu Tsai Neurosciences Institute), and the Stanford Alzheimer’s Disease Research Center (P30 AG066515). A.C.Y was supported by a Siebel Scholarship. F.K., G.S., T.F., W.S.-S., and A.K., are a part of the CORSAAR study supported by the State of Saarland, the Saarland University, and the Rolf M. Schwiete Stiftung.

## Author contributions

A.C.Y., F.K., A.K. and T.W.-C. conceptualized the study. M.W.M., N.L., I.C., W.S.-S., N.S., D.C., D.B., and A.C.Y. provided and organized tissue samples. A.C.Y. performed tissue dissociations. A.C.Y., N.S., D.P.L., R.T.V., D.G., and K.C. prepared libraries for sequencing. F.K., G.S., T.F., and A.C.Y. performed computational analysis, with F.K. leading advanced analysis and data management. P.M.L. developed the searchable web interface (Shiny app). W.S.-S. performed immunohistochemical stains. A.C.Y. and C.A.M. assembled figures. A.C.Y. wrote the manuscript with input from all authors. T.W.-C. and A.K. supervised the study.

## Competing interests

T.W.-C. is a founder and scientific advisor of Alkahest Inc.

Correspondence and requests for materials should be addressed to A.K. or T.W.-C.

## Methods

### Isolation of nuclei from frozen post-mortem medial frontal gyrus

Frozen medial frontal cortex tissue from post-mortem control and COVID-19 patients were obtained from the Stanford/ VA/ NIA Aging Clinical Research Center (ACRC) and the Saarland University Hospital Institute for Neuropathology, with approval from local ethics committees. Group characteristics are presented in Supplementary Table 1. The protocol for the isolation of nuclei was adapted from previous studies^32,34,35,148,149^, and performed in a BSL2+ Biosafety Cabinet wearing PPE. All procedures were carried out on ice or at 4°C. Briefly, 50 mg of postmortem brain tissue was dounce homogenized in 2 ml of Nuclei EZ Prep Lysis Buffer (Sigma, NUC101) spiked with 0.2 U μl^−1^ RNase inhibitor (Takara, 2313A) and EDTA-free Protease Inhibitor Cocktail (Roche, 11873580001) before incubating on ice for 5 min in a final volume of 5 ml. Homogenized tissue was filtered through a 100-μm cell strainer (Falcon, 352360), mixed with an equal volume of 50% iodixanol density gradient medium in PBS (OptiPrep, Sigma-Aldrich, D1556) to make a final concentration of 25% iodixanol. 30% iodixanol was layered underneath the 25% mixture. Similarly, 40% iodixanol was layered underneath the 30% iodixanol. In a swinging-bucket centrifuge, nuclei were centrifuged for 20 min at 3,000 r.c.f. After centrifugation, the nuclei were present at the interface of the 30% and 40% iodixanol solutions. Isolated nuclei were resuspended in 1% BSA with 0.2 U μl^−1^ RNase inhibitor, filtered twice through a 40 μm strainer (Flowmi), and counted on an TC20 automated cell counter (Bio-Rad) after the addition of Trypan Blue. We did not use statistical methods to pre-determine sample sizes, but our sample sizes are similar to those reported in previous publications^30–32^.

### Isolation of nuclei from frozen post-mortem choroid plexus

Frozen choroid plexus tissue was extracted from the lateral ventricles of post-mortem tissue obtained from the Stanford University Pathology department and the Saarland University Hospital Institute for Neuropathology, with approval from local ethics committees. Group characteristics are presented in Supplementary Table 1. All procedures were carried out on ice or at 4°C, and in a BSL2+ Biosafety Cabinet wearing PPE. Dounce homogenization or enzymatic dissociation successfully used for mice choroid plexi^29^ resulted in loss of nuclei integrity and low nuclei complexity (<50 median genes/ nuclei). We hypothesized that like shaking an apple tree, gentle pipetting of choroid plexi tissue in lysis buffer could liberate nuclei without needing to physically disintegrate the fibrous choroid matrix—and thus avoid collateral physical damage to nuclei. Specifically, 40 mg of choroid plexus tissue was thawed in 250 μl of 1% BSA with 0.2 U μl^−1^ RNase inhibitor until the tissue settled. 5 ml of lysis buffer (10 mM Tris, 10 mM NaCl, 3 mM MgCl2, 0.1% Nonidet P40 substitute (Roche/ Sigma, 11754599001), 0.2 U μl^−1^ RNase inhibitor, and protease inhibitor) was added and tissue incubated on ice for 10 minutes with gentle swirling every 2 minutes. 5 ml of 1% BSA was added and the tissue triturated 10 times with a 5 ml serological pipette. After centrifugation (500g, 5 min), pelleted nuclei were resuspended in 1% BSA with 0.2 U μl^−1^ RNase inhibitor; gently triturated 10 times with a 1 ml regular-bore pipette tip; and filtered twice through a 70 μm and then a 40 μm strainer (Flowmi). Debris was inspected on a brightfield microscope and nuclei were counted on an TC20 automated cell counter (Bio-Rad) after the addition of Trypan Blue.

### Droplet-based snRNA-sequencing

For droplet-based snRNA-seq, libraries were prepared using the Chromium Next GEM Single Cell 3’ v3.1 according to the manufacturer’s protocol (10x Genomics), targeting 10,000 nuclei per sample after counting with a TC20 Automated Cell Counter (Bio-Rad). 13 cycles were applied to brain parenchyma samples to generate cDNA, and 15 for choroid plexus samples. All samples underwent 15-16 cycles for final library generation. Generated snRNA-seq libraries were sequenced across two S4 lanes on a NovaSeq 6000 (150 cycles, Novogene).

### snRNA-seq quality control

Gene counts were obtained by aligning reads to the hg38 genome (refdata-gex-GRCh38-2020-A) using CellRanger software (v.4.0.0) (10x Genomics). To account for unspliced nuclear transcripts, reads mapping to pre-mRNA were counted. As previously published, a cut-off value of 200 unique molecular identifiers (UMIs) was used to select single nuclei for further analysis^34^. As initial reference, the entire dataset was projected onto two-dimensional space using Uniform Manifold Approximation and Projection (UMAP) on the top 20 principal components^150^. Three approaches were combined for quality control: (1) ambient cell free mRNA contamination was removed using SoupX^151^ for each individual sample; (2) outliers with a high ratio of mitochondrial (>5%, <200 features) relative to endogenous RNAs and homotypic doublets (> 5000 features) were removed in Seurat^152^; and (3) after scTransform normalization and integration, doublets and multiplets were filtered out using DoubletFinder with subsequent manual inspection and filtering based on cell type-specific marker genes^153^. Genes detected in fewer than 4 cells were excluded. After applying these filtering steps, the dataset contained 47,678 high-quality nuclei.

### Cell annotations

Seurat’s Integration function was used to align data with default settings. Genes were projected into principal component (PC) space using the principal component analysis (RunPCA). The first 80 (for global object), 30 (choroid plexus), or 25 (specific cell types) dimensions were used as inputs into Seurat’s FindNeighbors, FindClusters (at 0.2 resolution) and RunUMAP functions. Briefly, a shared-nearest-neighbor graph was constructed based on the Euclidean distance metric in PC space, and cells were clustered using the Louvain method. RunUMAP functions with default settings was used to calculate 2-dimensional UMAP coordinates and search for distinct cell populations. Positive differential expression of each cluster against all other clusters (MAST) was used to identify marker genes for each cluster^154^. We annotated cell-types using previously published marker genes^31,33–35^.

### Differential gene expression and sub-cluster analysis

Differential gene expression of genes comparing controls, COVID-19, and influenza samples— or comparing cell type sub-cluster markers—was done using the MAST^154^ algorithm, which implements a two-part hurdle model. Seurat default Log (fold change) > 0.25 (absolute value), adjusted *P* value (Bonferroni correction) < 0.05, and expression in greater than 10% of cells were required to consider a gene differentially expressed. Biological pathway and gene ontology enrichment analysis was performed using Enrichr^155^, Metascape^156^, or GeneTrail 3^157^ with input species set to *Homo sapiens*^156^ and using standard parameters. Docking, processing, and viral defense genes relevant to SARS-CoV-2 were chosen based on Iadecola, et al^16^. To identify microglia sub-cluster markers, differential expression analysis of cells grouped in each subcluster was performed against the remaining cells within the given cell-type. Markers were defined based on the MAST algorithm using only positive values with Log (fold change) > 0.25 (absolute value), adjusted P value (Bonferroni correction) < 0.01. Enrichment/ overrepresentation of the overlap between markers defining the COVID-19 microglia 2 cluster and the Mathys^34^ Alzheimer’s disease Mic1 cluster followed the hypergeometric probability, using the set of 17,926 protein-coding genes as background.

### Monocle trajectory analysis

Monocle3 (v.0.2.1.) was used to generate the pseudotime trajectory analysis in microglia^64^. Cells were re-clustered as above and used as input into Monocle to infer cluster and lineage relationships within a given cell type. Specifically, UMAP embeddings and cell sub-clusters generated from Seurat were converted to a *cell_data_set* object using SeuratWrappers (v.0.2.0) and then used as input to perform trajectory graph learning and pseudo-time measurement through reversed graph embedding with Monocle.

### Viral transcripts

To search for SARS-CoV-2 reads, raw fastq files were subjected to read alignment via Viral-Track^158^ or centrifuge^159^ using the human (GRCh38) genome reference. For Viral-Track both a collection of 12,163 consensus virus sequences from Virusite^160^ (release 2020.3) and 17,133 curated SARS-CoV-2 genomes from NCBI (downloaded on 29-09-2020) were used. For centrifuge, a preprocessed virus index compiled by genexa containing among other viruses 138 SARS-CoV-2 genomes was used. We also adopted a complementary approach^161^ focusing on SARS-CoV-2 reads, whereby barcoded but unmapped BAM reads were aligned using STAR to the SARS-CoV-2 reference genome, with a less stringent mapping parameter (outFilterMatchNmin 25-30) than the original Viral-Track pipeline.

### Cell-cell communication

Cell-cell interactions based on the expression of known ligand-receptor pairs in different cell types were inferred using CellChatDB^61^ (v.0.02). To identify potential cell-cell communication networks perturbed or induced in COVID-19 patient brains, we focused on differentially expressed ligands in choroid plexus epithelium, microglia, and astrocytes; and differentially expressed receptor pairs in excitatory and inhibitory neurons. Briefly, we followed the official workflow and loaded the normalized counts into CellChat and applied the preprocessing functions *identifyOverExpressedGenes, identifyOverExpressedInteractions*, and *projectData* with standard parameters set. As database we selected the *Secreted Signaling* pathways and used the pre-compiled human Protein-Protein-Interactions as a priori network information. For the main analyses the core functions *computeCommunProb, computeCommunProbPathway*, and *aggregateNet* were applied using standard parameters and fixed randomization seeds. Finally, to determine the senders and receivers in the network the function *netAnalysis_signalingRole* was applied on the *netP* data slot.

### GWAS

From the GWAS catalog^126^, we obtained GWAS risk genes for neurological disorders [Alzheimer’s disease (AD), amyotrophic lateral sclerosis (ALS), brain aging, multiple system atrophy (MSA), multiple sclerosis (MS), Parkinson’s disease (PD), and narcolepsy], psychiatric disorders [Attention deficit hyperactivity disorder (ADHD), autism, bipolar disorder, depression, psychosis, post-traumatic stress disorder (PTSD), and schizophrenia], and neurobehavior traits [Anxiety, suicidality, insomnia, neuroticism, risk behavior, intelligence, and cognitive function]. We removed gene duplicates and GWAS loci either not reported or in intergenic regions and used a *P* < 9 × 10^−6^ to identify significant associations^32^. Then, since GWAS signals can point to multiple candidate genes within the same locus, we focused on the ‘Reported Gene(s)’ (genes reported as associated by the authors of each GWAS study). Disorders and traits exhibiting a significant number of genes also perturbed in COVID-19 patients are highlighted. Following gene symbol extraction, we curated the gene set by (1) removing unknown or outdated gene names using the HGNChelper package (v.0.8.6), (2) converting remaining Ensembl gene IDs to actual gene names using the packages ensembldb (v.2.10.0) and EnsDb.Hsapiens.v86 (v.2.99.0), and (3) removing any remaining duplicates. We then calculated the overlap between each set of GWAS genes with the cell type-specific DEGs. Finally, a statistical enrichment of each overlap against background was calculated using a hypergeometric test with the total background size set equal to the number of unique RNAs mapped in our data set (29,431).

### Comparison of DEGs in chronic CNS disease

We compiled cell type-specific differentially expressed genes (DEGs) reported in published datasets for Alzheimer’s Disease (AD)^34^, Autism spectrum disorder (ASD)^33^, Huntington’s disease (HD)^123^, and Multiple sclerosis (MS)^122^. Lists of gene symbols were curated using the aforementioned approach. COVID-19 DEGs that overlap with those found across the selected CNS diseases were called shared, whereas those not previously reported were called unique to COVID-19. Statistical significance calculations of over-representation in DEG overlaps are based on cumulative hypergeometric *P* values analogous to the procedure described above.

### PVCA and PCA analysis

Briefly, to conduct the Principal Variance Component Analysis (PVCA), we aggregated the SoupX corrected raw counts for each gene and each biological sample using the *aggregateData* function of the muscat package (v.1.2.1). The resulting matrix was normalized by dividing each feature of a sample by the total counts from that sample, multiplied by a scaling factor of 100,000 and scaling the result using the function log(x+1). As variables we considered the sample annotation fields “Sample-ID”, “Patient-ID”, “Sex”, “Brain-region”, “Disease”, “ageBin”, and “nNucleiBin”. Since the number of nuclei and age are numeric variables but PVCA is designed to support factors, we assigned the values into ordered bins, more specifically, into five half-open (left-closed) intervals of size 1,000 starting at 2,000 for the number of nuclei and five similarly defined intervals of size 10 starting at 51 for the age. We set the algorithm cut-off for the minimal variance out of the total variance being explained to be 95%. For each single annotation variable, or first higher-order combinations of such, at cut-off of 0.005 was applied to consider them explanatory. All variables (or combinations of such) not passing the threshold were summarized as *Other* in the analysis. The residual is then defined as the remaining proportion of variance not being associated with any of the variables that are explanatory nor informative to a minor proportion. To conduct Principal Components Analysis (PCA) we aggregated the log-normalized cell counts from Seurat for each gene and sample using the *aggregateData* function from muscat and centered the gene expression vectors before computing eigenvectors.

### Overall in-silico analysis

Analysis and dissection of the data was performed using the statistical programming language R (v.3.6.3) using the following general-purpose package for loading, saving, and manipulating data, as well as generating plots, and fitting statistical models: dplyr (v.1.0.0), ggplot2 (v.3.2.2.), patchwork (v.1.0.1), openxlsx (v.4.1.5), bioconductor-scater (v.1.14.6), bioconductor-dropletutils (v1.6.1), bioconductor-complexheatmap (v.2.2.0), tidyverse (v.1.3.0), and lsa (v.0.73.2). All other tasks were performed on an x86_64-based Ubuntu (4.15.0-55-generic kernel) server cluster.

### Immunohistochemistry

Paraffin-embedded human brain tissue (medial frontal cortex, meninges, and choroid plexus) adjacent to tissue processed for snRNA-seq was subjected to immunohistochemistry (IHC). After deparaffinization and rehydration of 1-3 μm sections, peroxidases were blocked by incubation in 1% H2O2 for 15 minutes at room temperature. Heat antigen retrieval was performed by steaming at 98°C in target retrieval solution pH 6.1 (Dako, Carpinteira, CA, USA #S1699) for 30 minutes. Sections were allowed to cool down at room temperature. Following antigen retrieval, sections were incubated for 45 minutes at room temperature with the anti-SARS spike glycoprotein antibody 3A2 (rabbit, Abcam ab272420 1:100 diluted in Dako REAL antibody diluent #S2022), which has been validated in previous publications^20,41^. After three washes with wash buffer (Dako #S3006), the Dako REAL EnVision HRP kit (#K5007) was used for the visualization of the antibody reaction according to the manufacturer’s instructions. Sections were counterstained with Mayer’s hemalum (Sigma-Aldrich #1.09249). After dehydration, coverslips were mounted with Entellan (Merck #1.07961). Images were acquired with an Olympus BX 40 microscope, equipped with an Olympus SC30 digital microscope camera using the Olympus cellSens software.

## Data Availability

Raw sequencing data is deposited under NCBI GEO: GSE159812. Normalized counts data also available for download at: https://twc-stanford.shinyapps.io/scRNA_Brain_COVID19.

## Extended Data Figures

**Extended Data Figure 1.**
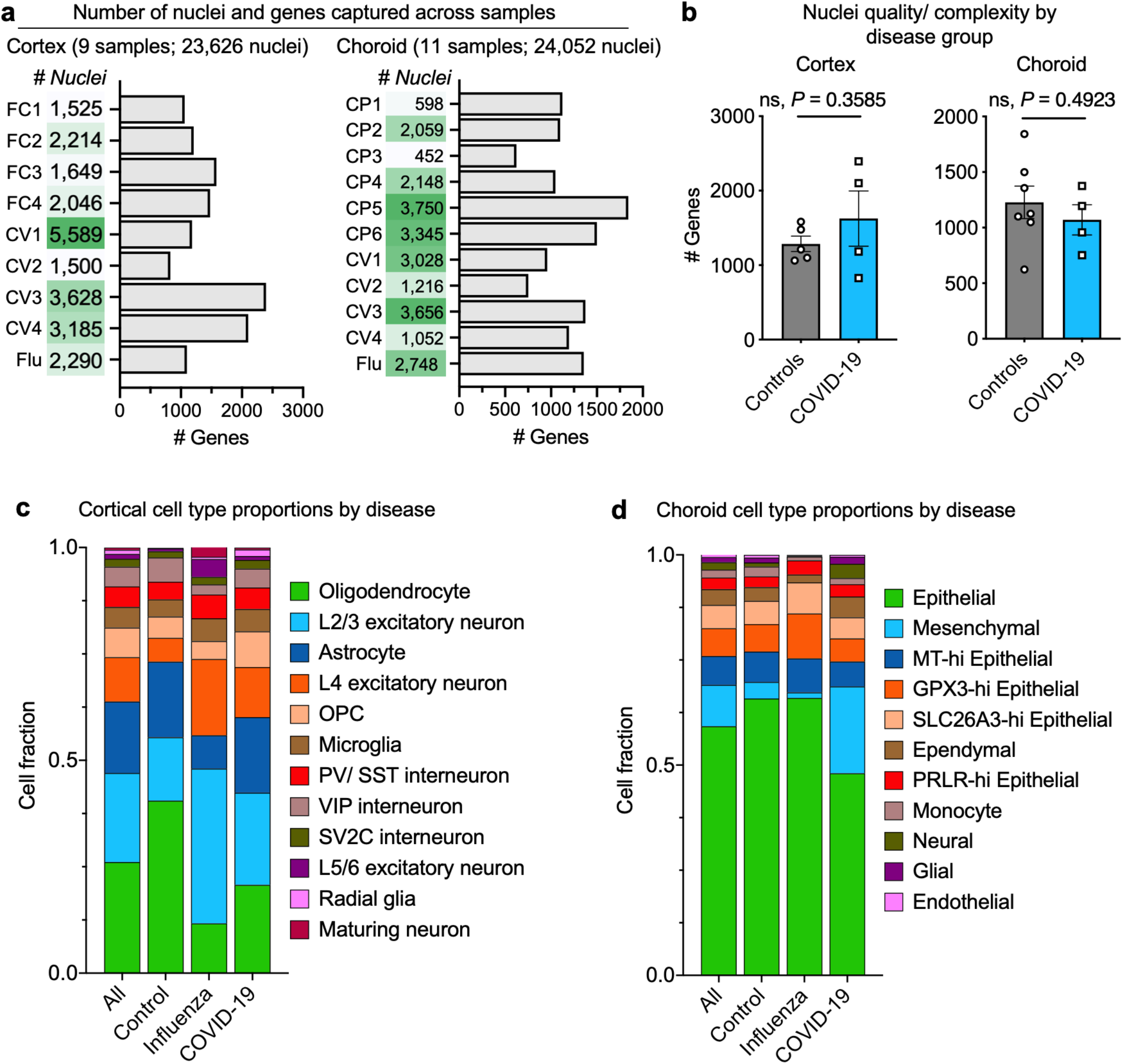
Characterization of human cortical and choroid plexi nuclei sequenced. **a**, Total number of nuclei and median number of genes of each human sample sequenced in medial frontal cortex and choroid plexus. **b**, Quantification of the median number of genes detected per nuclei in controls (non-viral and influenza) and COVID-19 samples in medial frontal cortex and choroid plexus (*n*=5-7 controls and *n*=4 COVID-19, two-sided t-test; mean +/− s.e.m.). **c**, **d**, Bar graph presenting frequency of nuclei for non-viral controls, influenza, and COVID-19 medial frontal cortex (**c**) and choroid plexus (**d**) sample groups.

**Extended Data Figure 2.**
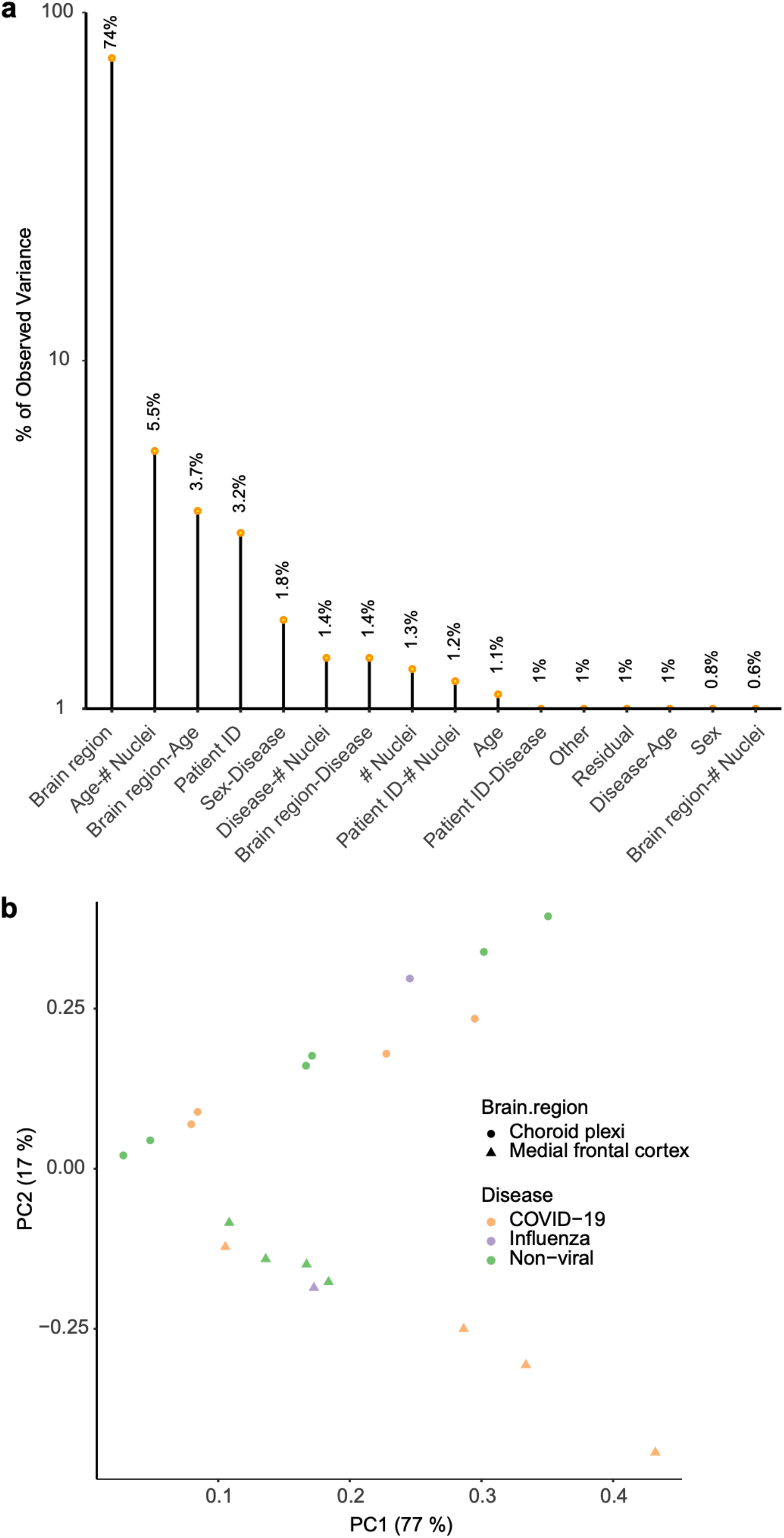
Gene expression variance analysis. **a**, Principal variance component analysis, displaying the gene expression variance explained by residuals (biological and technical noise) or experimental factors such as brain region, age, sex, and respective combinations. *n* = 15 samples. **b**, Principal component analysis (PCA) visualization of all samples, based on unscaled counts.

**Extended Data Figure 3.**
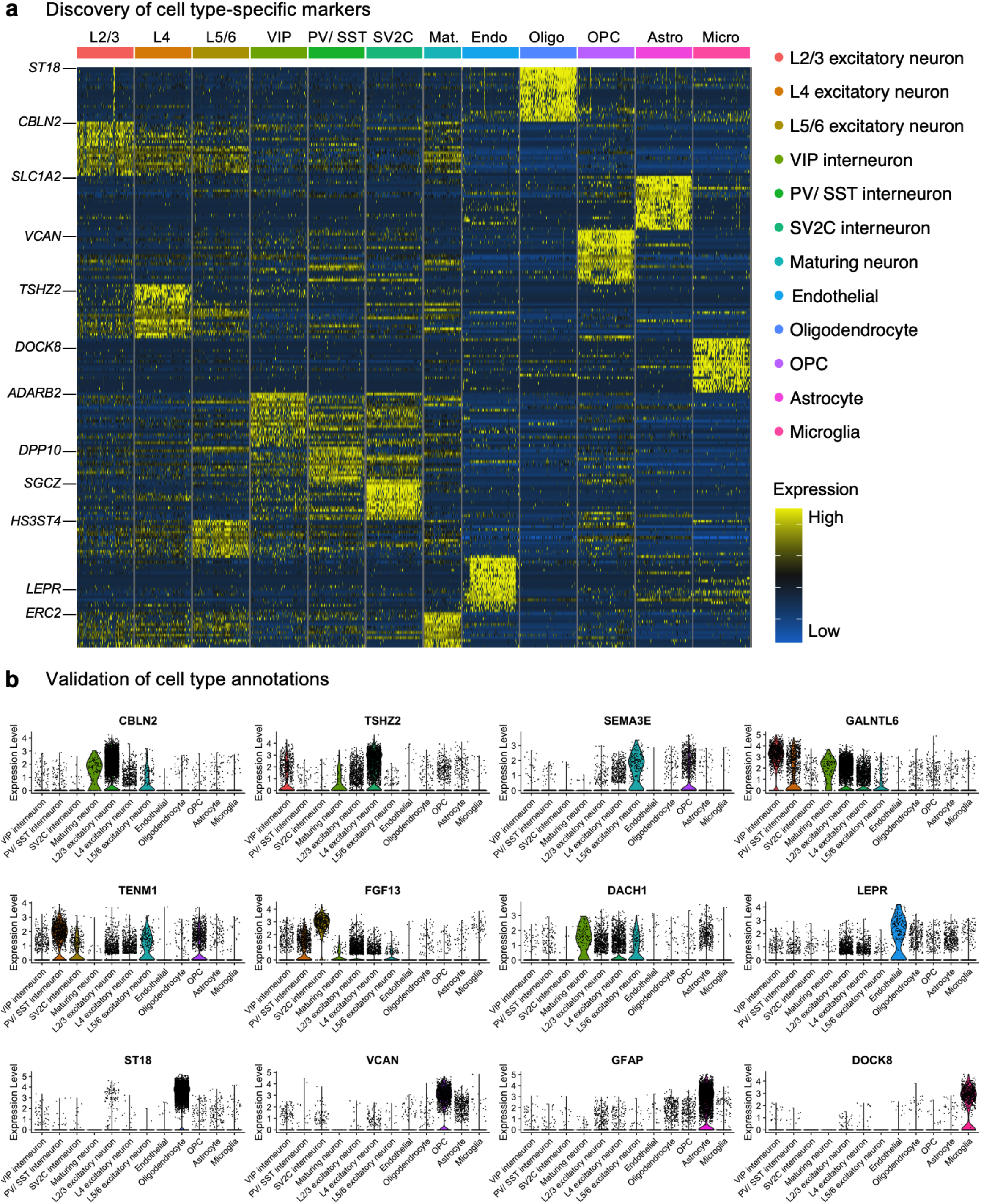
Markers for annotating parenchymal cells in the human frontal medial cortex. **a**, Discovery of the top cell type-specific genes across the 12 classes of cells captured in the human cortex. The color bar indicates gene expression from low (blue) to high (yellow). **b**, Validation of cell type annotations using established cell type-specific markers. Violin plots are centered around the median, with their shape representing cell distribution.

**Extended Data Figure 4.**
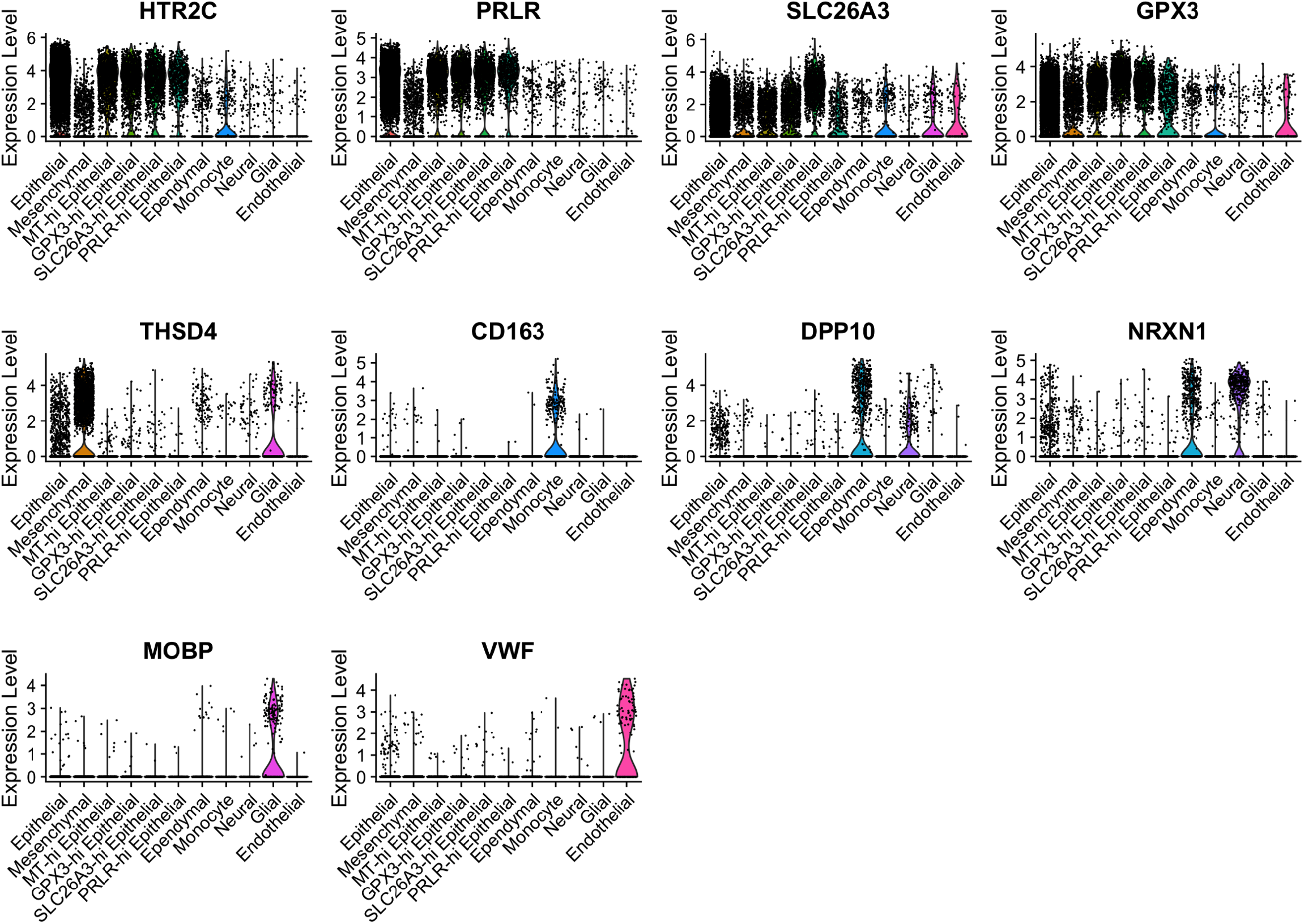
Markers for annotating parenchymal cells in the human choroid plexus. Validation of cell type annotations using established cell type-specific markers. Subsets of epithelial cells are marked by high expression of specific markers not previously described in mice. Violin plots are centered around the median, with their shape representing cell distribution.

**Extended Data Figure 5.**
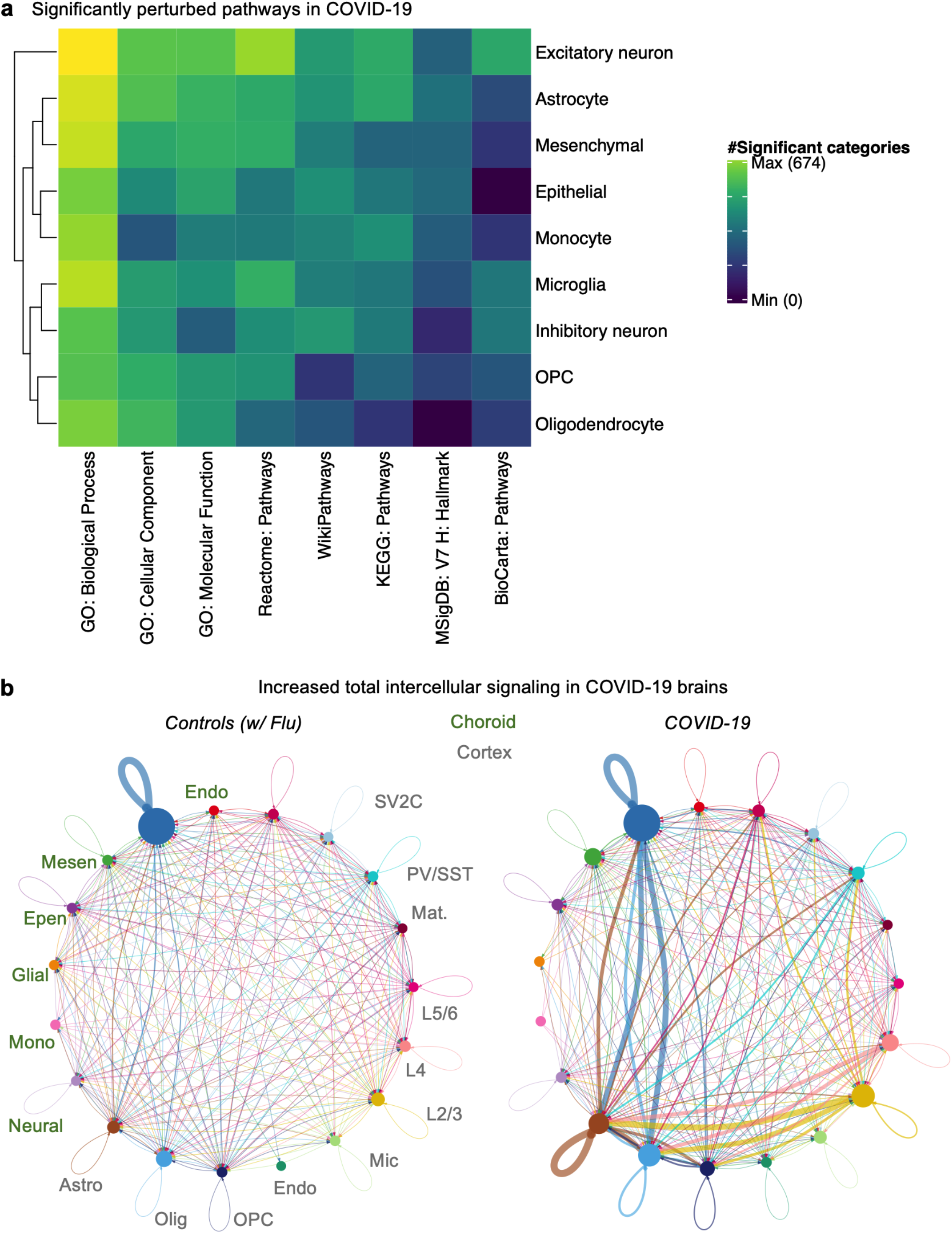
Cell-type specific changes in gene expression and intercellular signaling in the COVID-19 brain. **a**, Heatmap displaying the number of significant biological pathways among the set of differentially expressed genes in each cell type (FDR < 0.05, Fisher’s exact test with Benjamini-Hochberg correction). Number of significant pathways is indicated in graded black (high) to yellow (high). **b**, Circle plot showing the total inferred intercellular signaling network, aggregating across the signaling molecule interaction (i.e., ligand-receptor) CellChatDB database^61^.

**Extended Data Figure 6.**
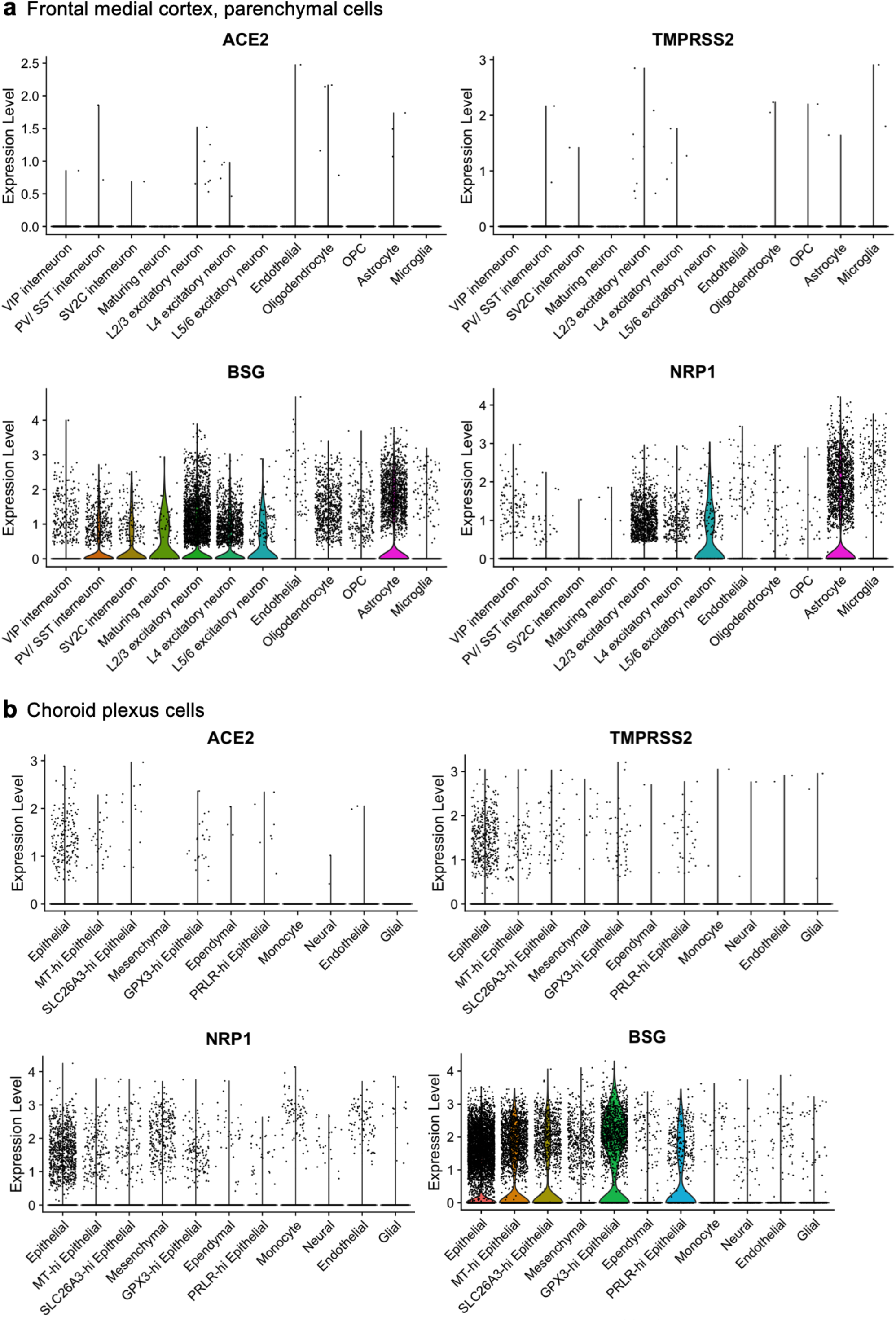
Expression of SARS-CoV-2 virus entry genes across cell types. **a**, **b**, Expression of SARS-CoV-2 entry receptors, established and putative, across cell types in the human medial frontal cortex (**a**) and choroid plexus (**b**). Violin plots are centered around the median, with their shape representing cell distribution.

**Extended Data Figure 7.**
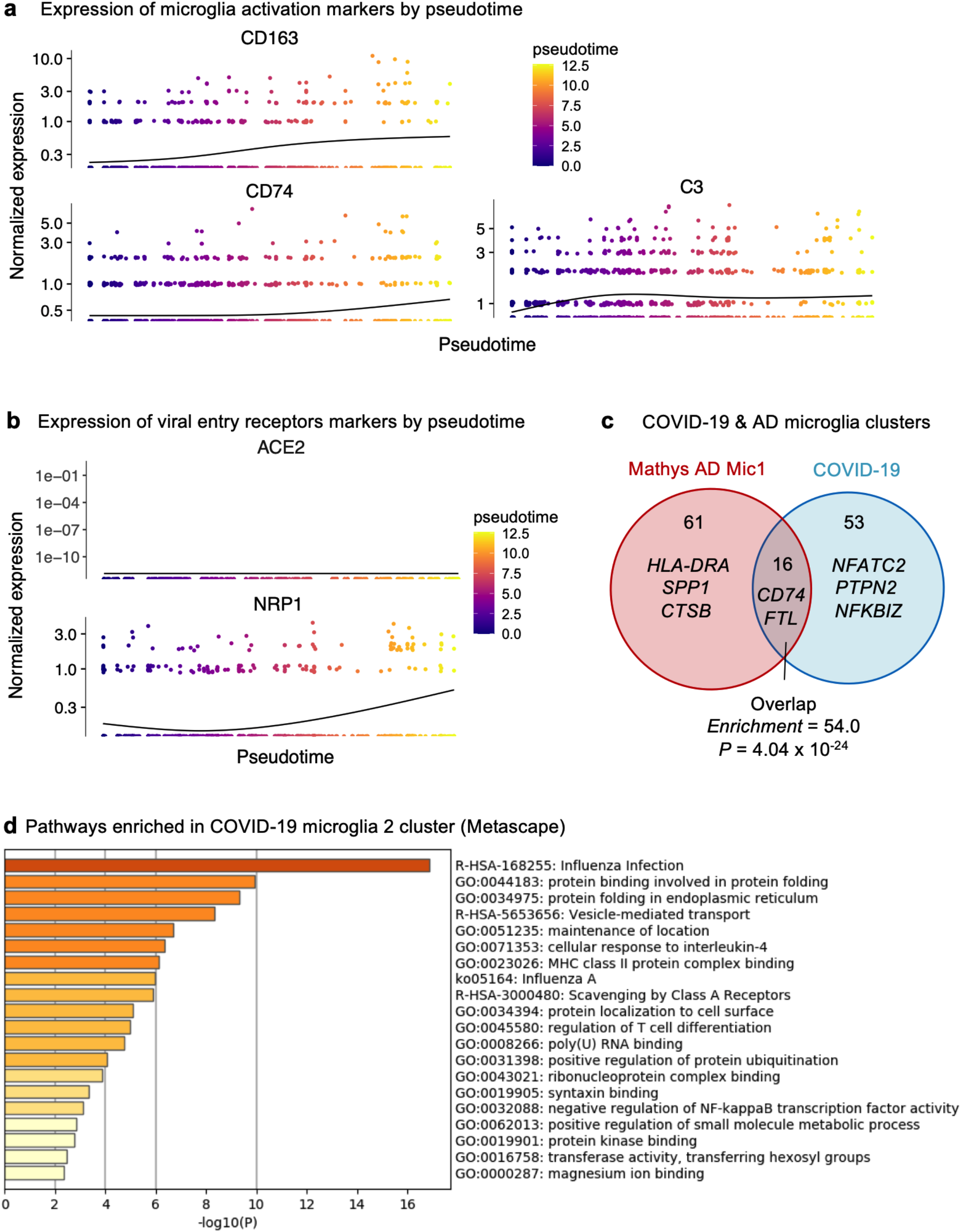
Evolution and properties of the COVID-19 enriched microglia cluster. **a**, **b**, Expression of established microglia activation markers (**b**) and SARS-CoV-2 entry receptors (**c**) along the pseudotime route in (**a**). Pseudotime is indicated in graded purple (high) to yellow (high). **c**, Overlap (hypergeometric test) between Alzheimer’s disease human activated microglia Mic1 marker genes^34^ and genes upregulated in the COVID-19 enriched Microglia 2 subcluster. **d**, Enriched biological pathways amongst Microglia 2 marker genes. Enrichment is based on FDR-corrected cumulative hypergeometric *P* values, with a *P* value-ranked gene-marker list (FDR < 0.01, log2(mean gene expression across Microglia 2/ mean gene expression across all microglia) > 0.25, MAST).

**Extended Data Figure 8.**
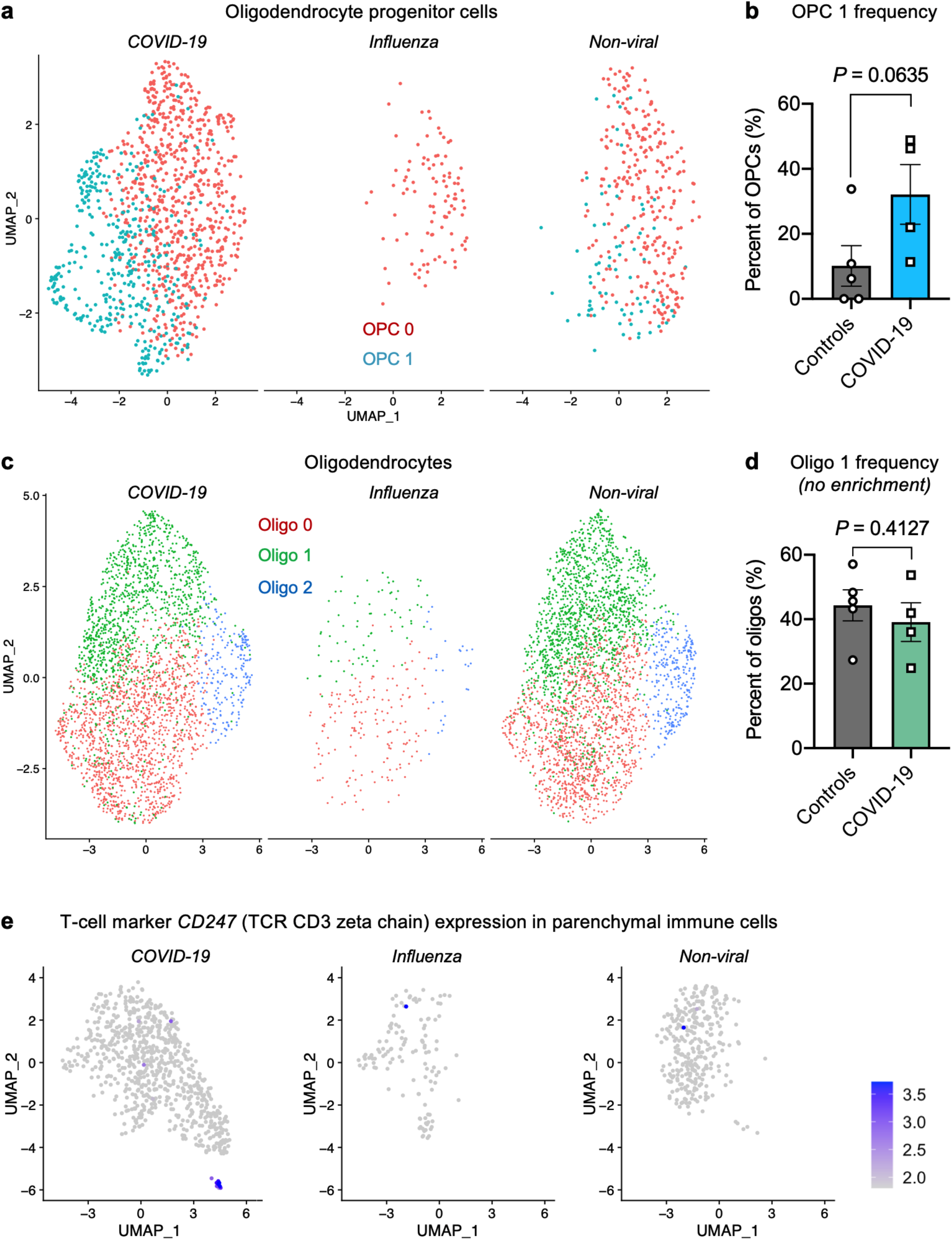
Evaluation of COVID-19 enriched subpopulations in other parenchymal glia. **a**, UMAP projection of oligodendrocyte progenitor cells (OPCs, *n* = 1,634 from 4 non-viral controls, 1 influenza and 4 COVID-19 individuals) and the emergence of COVID-19 enriched subcluster. **b**, Quantification of the frequency of the COVID-19 enriched OPC subcluster as a proportion of all OPCs (*n*=5 controls and *n*=4 COVID-19, two-sided t-test; mean +/− s.e.m.). **c**, **d**, As in (**a**-**b**) but for oligodendrocytes. **e**, Expression of the established T-cell marker *CD247* (encoding TCR CD3 zeta chain) among immune cells captured in human medial frontal cortex.

**Extended Data Figure 9.**
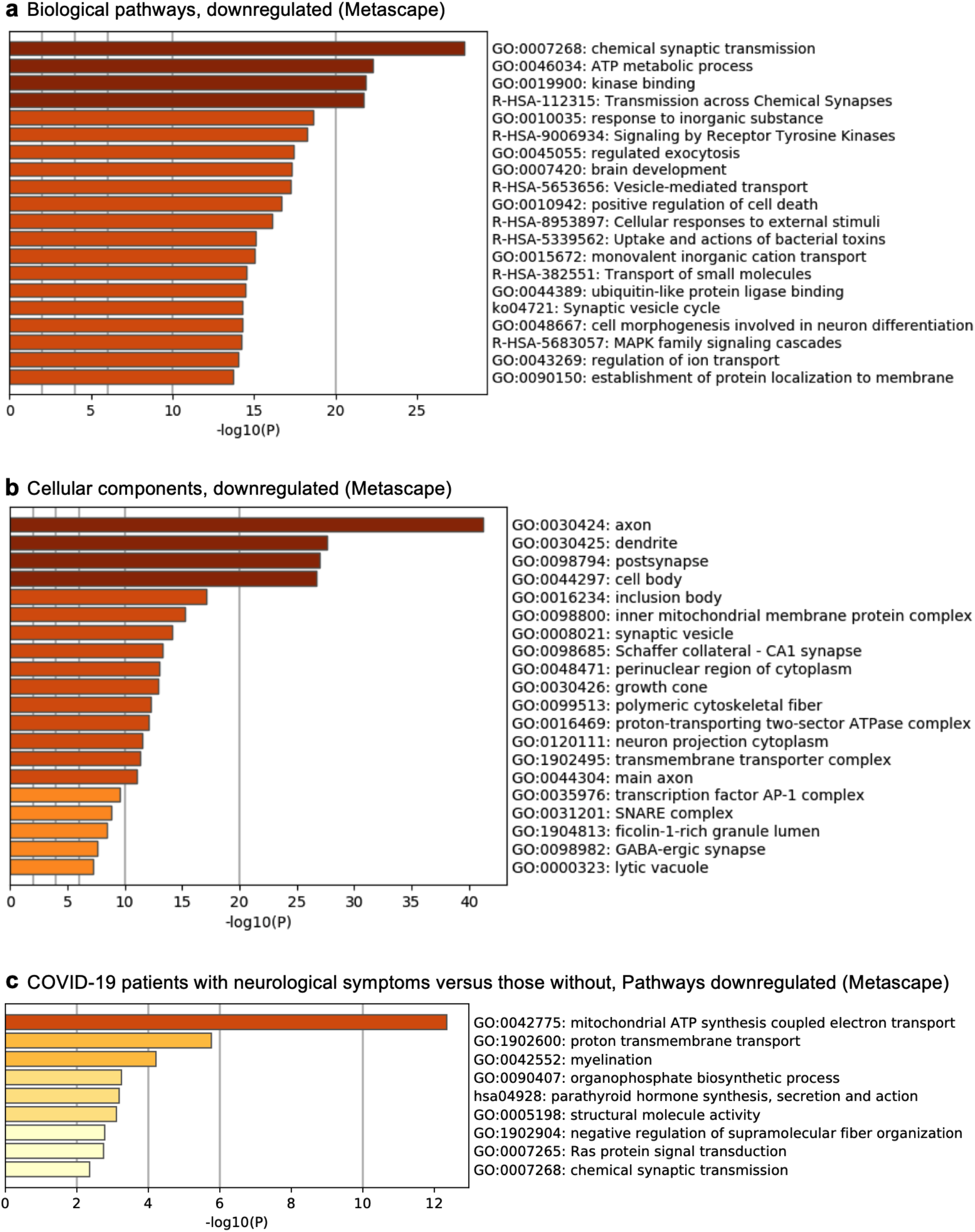
Processes downregulated in excitatory neurons in COVID-19. **a**, Enriched biological pathways amongst downregulated genes in COVID-19 excitatory neurons (*n* = 7,680 from 4 non-viral controls, 1 influenza and 4 COVID-19 individuals). Enrichment is based on FDR-corrected cumulative hypergeometric *P* values, with a P value-ranked gene-marker list (Log (fold change) > 0.25 (absolute value), adjusted *P* value (Bonferroni correction) < 0.05, MAST). **b**, As in (**a**) but for Gene Ontology cellular components. **c**, As in (**a**) but comparing COVID-19 patients with neurological symptoms versus those without.

**Extended Data Figure 10.**
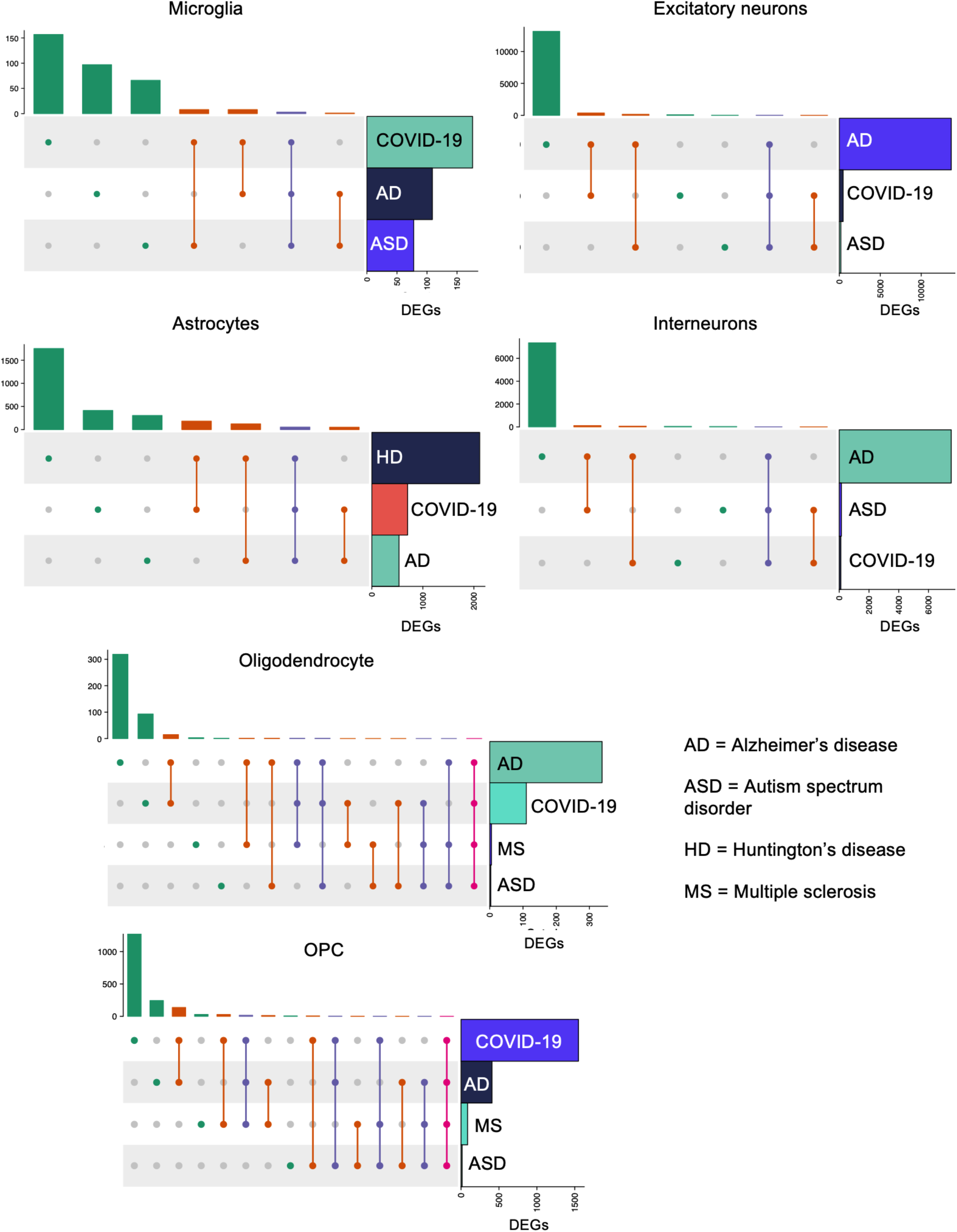
Comparison of cell type-specific changes in COVID-19 with chronic CNS diseases. Upset matrix plots comparing the cell type-specific genes expression changes in COVID-19 frontal cortex with those previously published for Alzheimer’s Disease (AD)^34^, Autism spectrum disorder (ASD)^33^, Huntington’s disease (HD)^123^, and Multiple sclerosis (MS)^122^.

